# Reserpine prolongs lifespan but compromises heat-stress resilience in *Drosophila melanogaster*

**DOI:** 10.1101/2025.08.20.671049

**Authors:** Vaibhav Tiwary, Nares Trakooljul, Shahaf Peleg

## Abstract

Pharmacological modulation of monoaminergic signaling, a process targeted by many therapeutic and recreational drugs via receptors, transporters, degradation enzymes, or reuptake mechanisms, is emerging as a promising aging intervention and as a strategy to treat various maladies. Monoamines (including dopamine, serotonin, and norepinephrine) are central to the regulation of mood, movement, sleep, memory, and systemic physiology. Here, we demonstrate that Reserpine, chronic inhibitor of the vesicular monoamine transporter (VMAT), robustly extends lifespan in *Drosophila melanogaster* in a dose-dependent manner. However, reserpine-treated flies also exhibit reduced locomotor activity and impaired survival under acute heat stress, indicating a context-dependent trade-off between lifespan extension and stress resilience. Transcriptomic profiling revealed that reserpine induces a transcriptionally repressed, low-energy state characterized by downregulation of metabolic, immune, and stress-response genes in treated aged animals. Notably, under heat stress, reserpine blunts the induction of canonical protective genes, including heat shock proteins and antioxidant genes, resulting in increased proteotoxic vulnerability. These findings highlight the potential trade-offs of monoaminergic modulation and support further investigation of VMAT inhibitors, monoamine modulators and other hypertension drugs as geroprotective agents.

## Introduction

Aging is a multifactorial biological process that is characterized by the progressive loss of physiological integrity, functional decline and increased vulnerability to stress and disease. It is orchestrated by an intricate interplay of genetic programs, metabolic states, and environmental exposures^1^. These pathways play central roles in nutrient sensing, metabolic regulation, neurotransmission, and stress response, while their manipulation has been shown to extend lifespan and delay age-related decline across multiple model organisms ^1–4^.

A substantial body of research has established that both lifespan and health span can be modulated by targeting evolutionarily conserved signaling pathways, notably the insulin/IGF-1, mTOR and AMPK pathways, as well as monoaminergic signaling ^1–5^. For example, change in monoamine concentrations due to overactivation of Monoamine oxidases (MAO) has been found to play a determining role in aging associated pathologies like neurodegeneration, pulmonary diseases, cancer and metabolic disorders including obesity and diabetes ^6^. The discoveries of modulating signaling pathways for longevity has prompted efforts to identify pharmacological agents capable of safely mimicking the benefits of dietary restriction and stress resistance without requiring drastic lifestyle interventions ^7,8^, including repurposing existing FDA approved drugs for anti-aging therapies ^9,10^.

Notably, several clinically approved antihypertensive drugs that were initially developed for cardiovascular control have emerged as candidate geroprotectors. These include, rilmenidine (an I₁-imidazoline receptor agonist), metolazone (a thiazide-like diuretic) and reserpine (a vesicular monoamine transporter inhibitor). In *Caenorhabditis elegans*, each of these three drugs has been shown to extend lifespan and enhance markers of health span, such as locomotion and thermotolerance ^11–13^. The convergence of pharmacological action between these antihypertensive compounds and well-characterized longevity interventions underscores the central role of neuroendocrine and metabolic adaptation in aging. However, while promising, these mechanisms remain incompletely understood, with respect to how they affect acute stress resilience, their secondary effects and how and if they can be translated to more complex organisms.

Recent studies in older humans show that long-term use of antihypertensive drugs, especially calcium channel blockers, is linked to slower biological aging and lower frailty. The study by Tang, et al. ^14^ found that antihypertensive drug use overall was associated with slower biological aging, specifically showing a decrease in DNA methylation age as measured by the PCGrimAge clock. When examining drug subcategories, calcium channel blockers (CCBs) consistently showed links to decreased DNA-methylation ages (PCHorvathAge; PCSkin & bloodAge; PCPhenoAge; PCGrimAge) and in functional biological ages (functional age index; frailty index) ^14^. This study population primarily consisted of individuals using medications for conditions such as hypertension, diabetes, and hyperlipidemia, indicating that they had the relevant disorders for which these drugs are prescribed. While antihypertensives clearly reduce cardiovascular risks and mortality, they do not always lead to slower epigenetic aging (DNAmAge), indicating that the relationship between these drugs and biological aging is complex and needs further investigation to understand the mechanisms involved ^15^.

Reserpine, a root extract of the perennial shrub of the Rauwolfia family was historically used in traditional medicine and later incorporated into modern pharmacology. More recently it was used as a therapy for reducing blood pressure but nowadays used as a second-line treatment ^13,16^. Reserpine is a VMAT-1 and VMAT-2 (vesicular monoamine transporter) inhibitor previously used as an antihypertensive and antipsychotic. By blocking VMAT in neurons, dopamine (DA), serotonin (5-HT) and octopamine are depleted from vesicles, thus reducing their release to the synapse and further to the post-synaptic neuron ^17^. VMAT blockade consequently mimics neurotransmitter-deficient states: for example, reserpine-treated cells accumulate cytosolic DA, generate reactive oxygen species (ROS), and trigger apoptosis ^18^. Notably, reserpine’s broad action differs from more selective monoamine drugs: for instance, the antipsychotic haloperidol blocks D2 receptors ^19^, whereas reserpine’s inhibition of VMAT results in elimination of multiple neurotransmitter stores such as depleting dopamine, serotonin, norepinephrine and histamine ^20^. As such, reserpine treatment might result in engagement of complex downstream pathways which could be resulting in extended lifespan.

In the early 1950s reserpine was introduced as a first-line antihypertensive and, for a short time, as an antipsychotic ^20^. Clinicians observed that reserpine often induced depression-like symptoms that typically abated after stopping the drug. These findings underpinned the mono-amine-depletion hypothesis, which posits that loss of catecholamines (dopamine, norepinephrine) causes depression and helped to justify the drug’s withdrawal from routine use ^20^. A 2023 systematic review Strawbridge, et al. ^20^ examined 35 human studies on reserpine and concluded that the link between mono-amine depletion and depression is inconsistent and poorly supported, urging a nuanced reinterpretation of the classic monoamine hypothesis rather than a simple causation model. Another pilot study by Siddiqui, et al. ^21^ suggests that reserpine could also be reconsidered for refractory hypertension ^22^.

Reserpine has been shown to extend lifespan in *Caenorhabditis elegans* through two distinct and parallel pathways ^3^. One pathway involves dopaminergic signaling via the D2-like receptor DOP-3, which acts through the G-protein GOA-1 to activate the transcription factor JUN-1, ultimately upregulating the ABC transporter MRP-1; a gene associated with detoxification and stress resistance. The second pathway involves ERI-1, a 3′–5′ exoribonuclease that suppresses RNA interference; its loss enhances neuronal sensitivity to RNAi and is also required for the lifespan-extending effects of reserpine. These mechanisms operate independently of canonical longevity pathways such as insulin/IGF-1 signaling or dietary restriction in *C.elegans* ^3^. While this study provides key mechanistic insights into how reserpine promotes longevity in C. elegans, it does not offer a broader systems-level view of how these pathways integrate and modulate with metabolic, immune, or stress-response networks, nor does it address whether reserpine can operate in more complex organisms for lifespan extension.

Notably, a key question in translational Geroscience is whether interventions that promote lifespan necessarily enhance stress resistance or whether trade-offs emerge between longevity and stress adaptability^23,24^. For example, our recent work in flies revealed that reducing the levels of chm, a lysine acetyltransferase, leads to life span extension in flies while impairing their capacity to survive starvation in cold temperatures ^25^. To bridge this gap, we employed the model organism Drosophila melanogaster to investigate the effects of chronic dietary reserpine supplementation. Many of the neurotransmitters utilized by mammals are also used by flies, such as the monoamines histamine, serotonin (5HT), and dopamine (DA)^26^. Since histamine is produced by the enzyme histidine decarboxylase and plays a part in the visual system, it has been thoroughly studied in insects ^27^. In Drosophila, octopamine is a major stress/arousal transmitter and analogue of norepinephrine ^28^ while dopamine and serotonin regulate arousal, feeding and mood ^29–31^. Overall, Drosophila possesses a conserved neuroendocrine architecture ^26^, robust stress-response pathways, and well-characterized aging phenotypes^32^, making it an ideal system to explore pharmacological interventions^33^ with potential relevance to human biology ^34^.

In this study, we assessed how reserpine affects lifespan, heat-stress resilience, and global gene expression patterns. By integrating survival assays and transcriptomic profiling, our study uncovers how pharmacological modulation of monoaminergic signaling induces a low-energy, longevity-associated state but at the cost of reduced locomotor activity and suppressed acute stress resilience such as heat stress. These results reveal a mechanistic trade-off with implications for both fundamental aging biology and the rational development of geroprotective therapies using FDA-approved drugs.

## Results

### Lifespan Extension by Chronic Dietary Reserpine Treatment in Male Drosophila

We first tested the impact of various concentrations of reserpine on the lifespan of male flies. The first pilot experiment was with various concentrations ranging from 0 to 800uM of drug in fly food (**Figure S1).** We noticed a detectable life span increase in reserpine-treated flies between 200 µM to 800 µM. We next increased the concentration of the drug to 1000uM along with a condition of start of drug treatment at around half the lifespan of fly with (0-800 µM; reserpine introduced at day 31) as 800µM was most promising in the first pilot. Starting the treatment in midlife with 800µM failed to increase the median lifespan, thus suggesting that early life treatment was crucial for lifespan extension. Furthermore, increasing the treatment to 1000 µM showed further increase lifespan compared with 800µM **(Figure S2)**.

We next focused on higher concentrations of 1000 µM and 1500 µM. Indeed, chronic dietary administration of higher dosage reserpine significantly extended the median, mean, and maximum lifespan (**Figure 1**). Specifically, Kaplan–Meier survival analysis demonstrated an increase in median survival from 53 days in controls to 57 days at 1000 µM and 58 days at 1500 µM reserpine (log-rank test: p < 0.0001 for all comparisons). The mean lifespan also increased from 51.5 days in controls to 54.9 days at 1000 µM and 56.7 days at 1500 µM. The maximum lifespan rose from 68 days in controls to 81 days at both 1000 µM and 1500 µM concentrations (**Table 1**).

**Figure 1.**
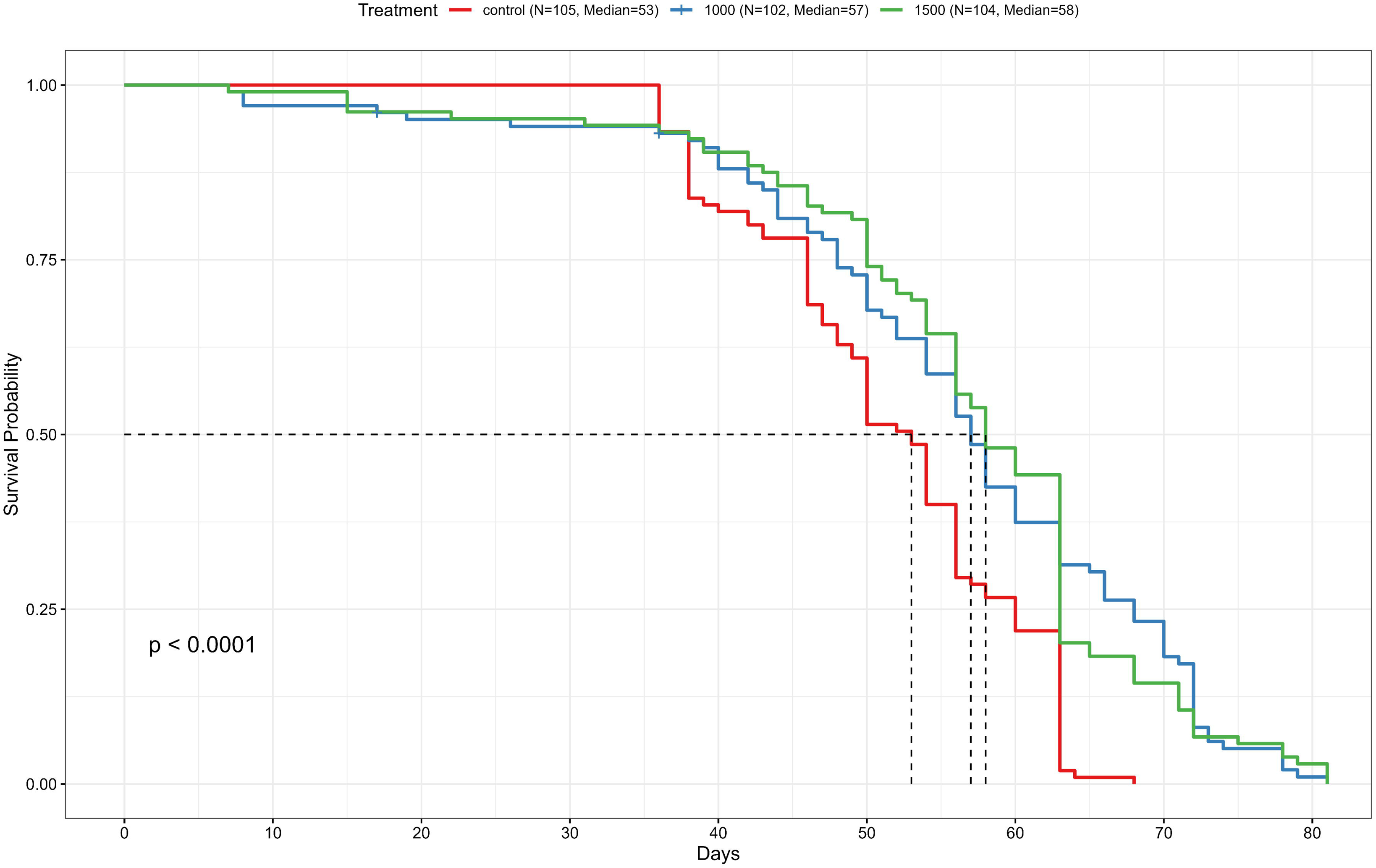
Reserpine treatment extends lifespan in male Drosophila melanogaster. Kaplan–Meier survival curves for male flies chronically treated with 0 (control), 1000, or 1500 µM reserpine under standard conditions. Median lifespans were 53 days for control (N = 105), 57 days for 1000 µM (N = 102), and 58 days for 1500 µM reserpine-treated flies (N = 104). Each treatment group included 3 independent biological replicates each containing an average of n ∼ 35 flies per vial. Overall group differences were statistically significant (log-rank test, p < 0.0001).

**Table 1.**
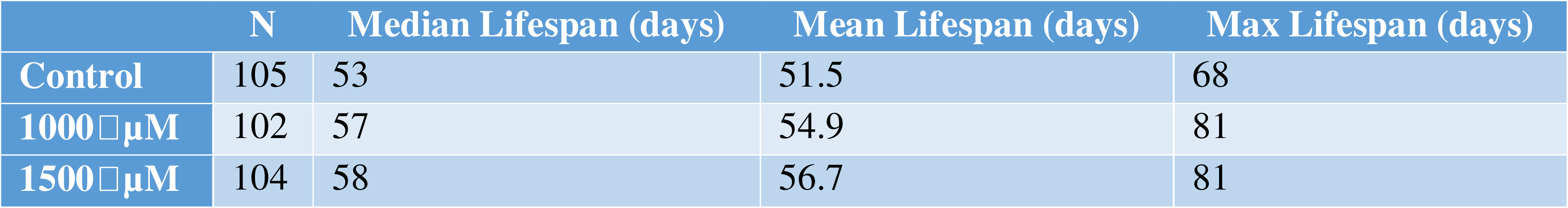
Summary of Median, Mean and Maximum Lifespan Extension under Reserpine Treatment.

|  | N | Median Lifespan (days) | Mean Lifespan (days) | Max Lifespan (days) |
| --- | --- | --- | --- | --- |
| Control | 105 | 53 | 51.5 | 68 |
| 1000 μM | 102 | 57 | 54.9 | 81 |
| 1500 μM | 104 | 58 | 56.7 | 81 |

### Reserpine reduces climbing ability and heat stress tolerance

Previous studies in *C.elegans* showed reserpine increased thermotolerance and locomotion ^13^. Contrarily, it was evident that reserpine treated flies exhibit decreased activity in survival assays. To quantify the fly activity, we conducted climbing assay ^35^. Reserpine-treated flies had severely reduced activity in climbing assay after 13 days of treatment. Tukey’s post hoc comparisons revealed a significant reduction in Weighted Climbing Index (WCI) in both the 1000 µM (mean = 1.52 ± SEM) and 1500 µM (mean = 1.43 ± SEM) groups compared to the control group (mean = 3.78 ± SEM. No significant difference was observed between the 1000 µM and 1500 µM groups (p = 0.9286). These results indicate that chronic treatment with the drug at both concentrations significantly impairs climbing ability in adult flies, with no additional impairment at the higher dose (**Figure 2**).

**Figure 2.**
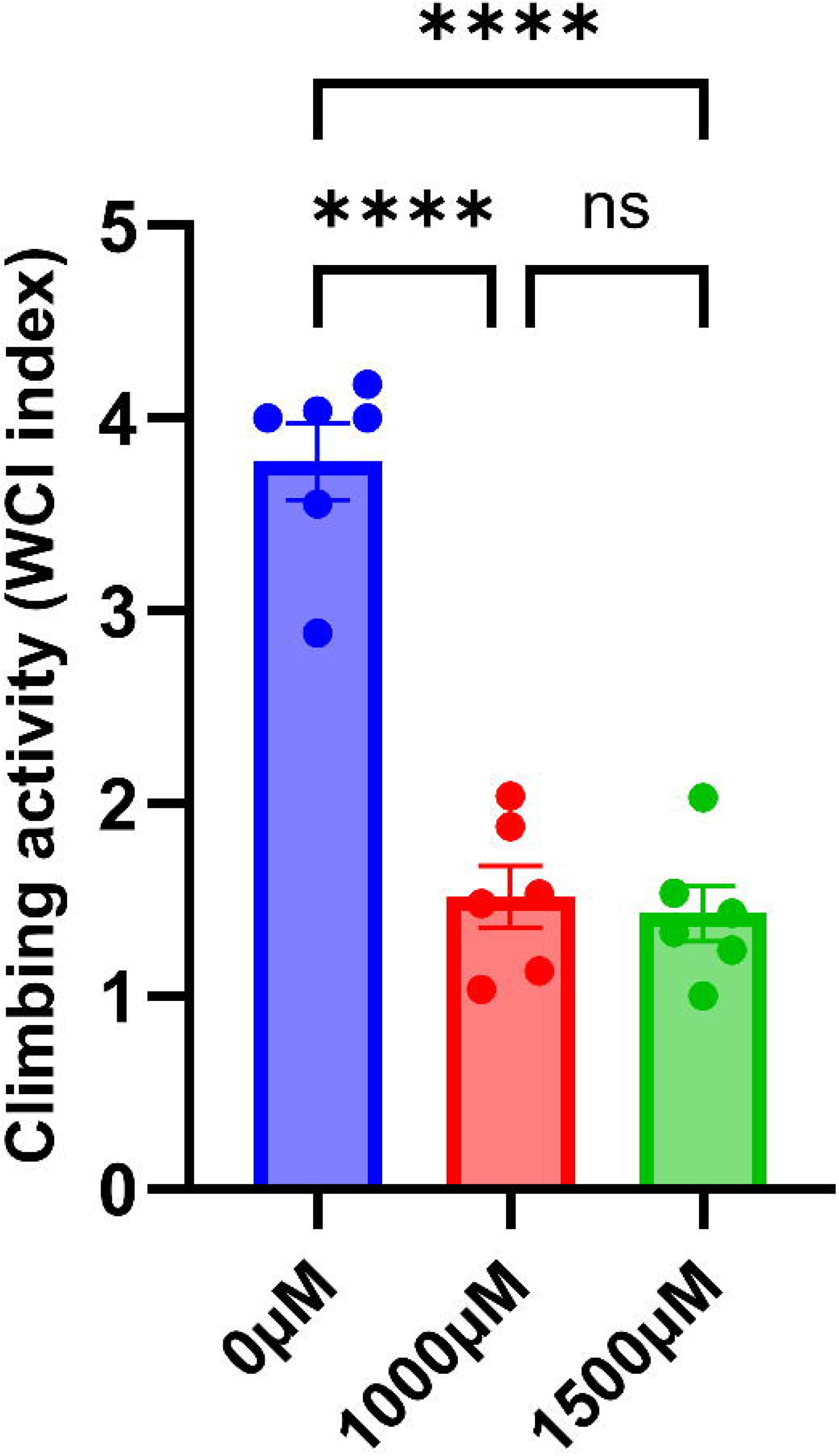
Chronic Reserpine treatment impairs climbing ability in Adult Drosophila. Flies were treated with 0 µM, 1000 µM, or 1500 µM of the test compound for 12 days, and climbing ability was assessed on day 13 using a negative geotaxis assay. Each bar represents the Weighted Climbing Index (WCI), calculated based on fly distribution across five equal vertical quadrants at 10 seconds post-stimulation. Each point represents one biological replicate (n = 6 per group; 25–31 flies per replicate). Bars show mean ± SEM. Statistical analysis was performed using one-way ANOVA with Tukey’s multiple comparisons test. ****p < 0.0001; ns = not significant.

We have recently shown that long lived fly mutant display reduced resilience to temperature induced stress ^36^. Therefore, we subjected young control and treated flies to dry heat stress after 12 days of treatment at standard conditions. Under continuous heat stress conditions at day 12 of their life (31 °C), reserpine significantly reduced survival in flies in a dose-dependent manner (**Figure 3**). Median survival decreased from 7 days in the control group (n = 115) to 5 days with 1000 µM reserpine (n = 115), and 4.5 days with 1500 µM reserpine (n = 120). Correspondingly, the mean lifespan declined from 7.06 days in controls to 5.97 days and 4.31 days in the 1000 µM and 1500 µM groups, respectively, while the maximum lifespan under the heat stress remained at 12 days across all groups. Log-rank analysis confirmed a highly significant difference between treatment groups (p < 0.0001). These results are consistent with study by Bressan, et al. ^37^ who found that reserpine impaired thermal tolerance, heat-avoidance behavior and locomotor activity in *Drosophila* ^37^. Together, the data suggest that while reserpine extends lifespan under normal conditions, it compromises the organism’s ability to withstand acute environmental stress and reduces activity.

**Figure 3.**
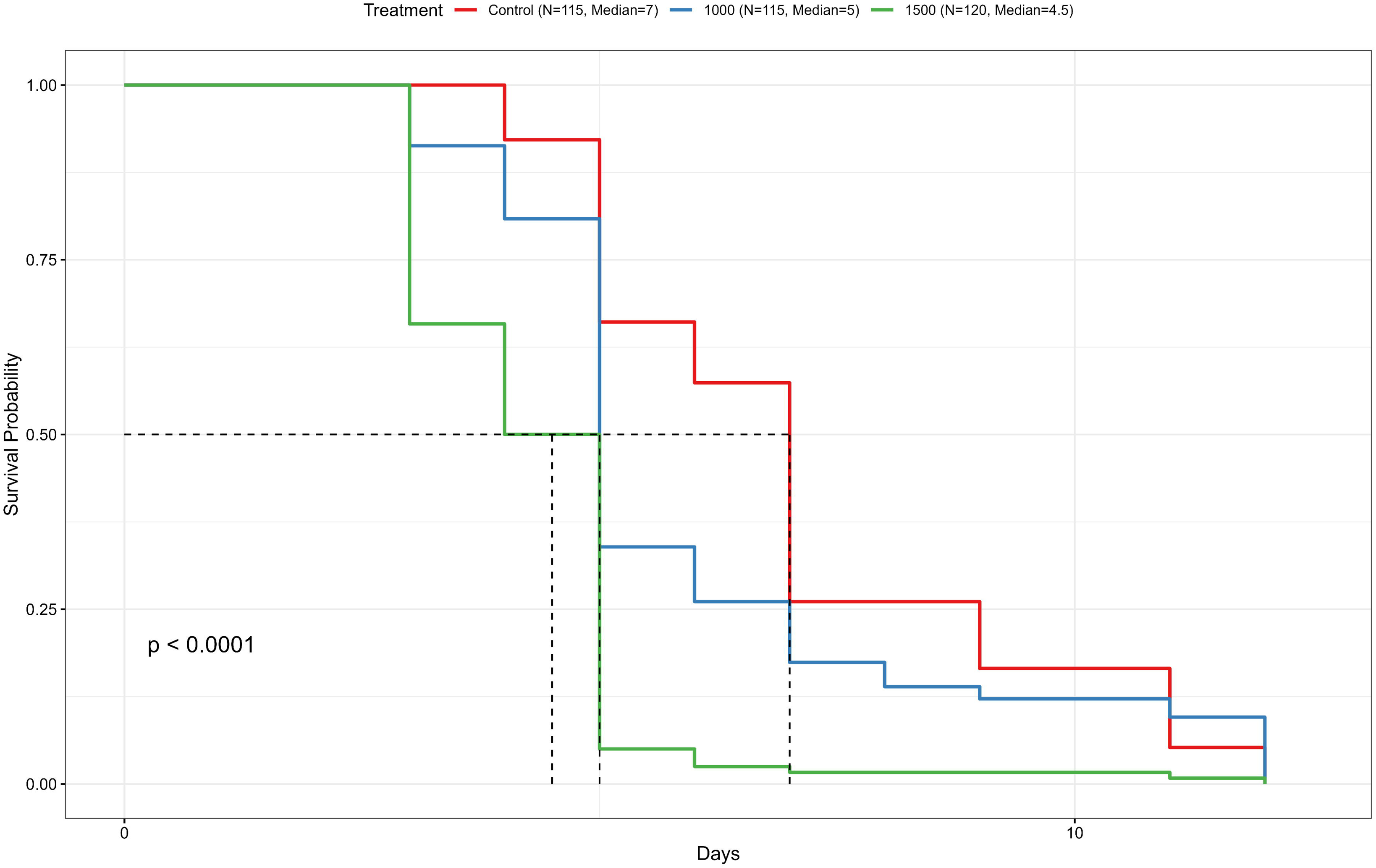
Reserpine treatment increases mortality under heat stress in young male flies. Kaplan–Meier survival analysis under continuous heat stress (31 °C) shows median survival times of 7 days (control), 5 days (1000 µM), and 4.5 days (1500 µM). Each treatment group included 3 independent biological replicates each containing an average of n ∼ 35 flies per vial. Overall group differences were significant (log-rank test, p < 0.0001).

### Reserpine Alters the Transcriptome of Aged Drosophila

Various studies have shown that are distinct transcriptomic and metabolic shifts occurs during aging ^38,39^ and Heat shock proteins (Hsps) are known modulators of stress-resistance and lifespan in Drosophila ^40^. Therefore, we first compared the transcriptome of 43 days control vs chronic reserpine treated male flies. RNA-seq analyses revealed that reserpine induces a robust, condition-specific transcriptomic reprogramming in aged Drosophila (**Figure 4**). Principal Component Analysis (PCA) demonstrated a clear separation between control and reserpine-treated samples, highlighting a substantial impact of reserpine on the aged transcriptome (**Figure 4A**). A heatmap of normalized expression values for differentially expressed genes (DEGs; padj < 0.05, n = 3964) showed distinct expression patterns and consistent clustering by treatment group (**Figure 4B**). MA plot highlights significantly changed genes (adjusted *p* < 0.05) with log2 fold change ±0.58. Many genes showed moderate yet significant fold changes, reflecting a subtle but coordinated transcriptomic reprogramming in response to reserpine (**Figure 4C**). The volcano plot summarizes statistical significance versus fold change. Genes such as *Gillspa92*, *spn1*, *GstD5*, *Cyp6g1*, and *lncRNA:CR43417* were significantly downregulated, whereas *Cyp28a5*, *Cyp6w1*, *CG2003*, and *Amyrel* were significantly upregulated (**Figure 4D**).

**Figure 4.**
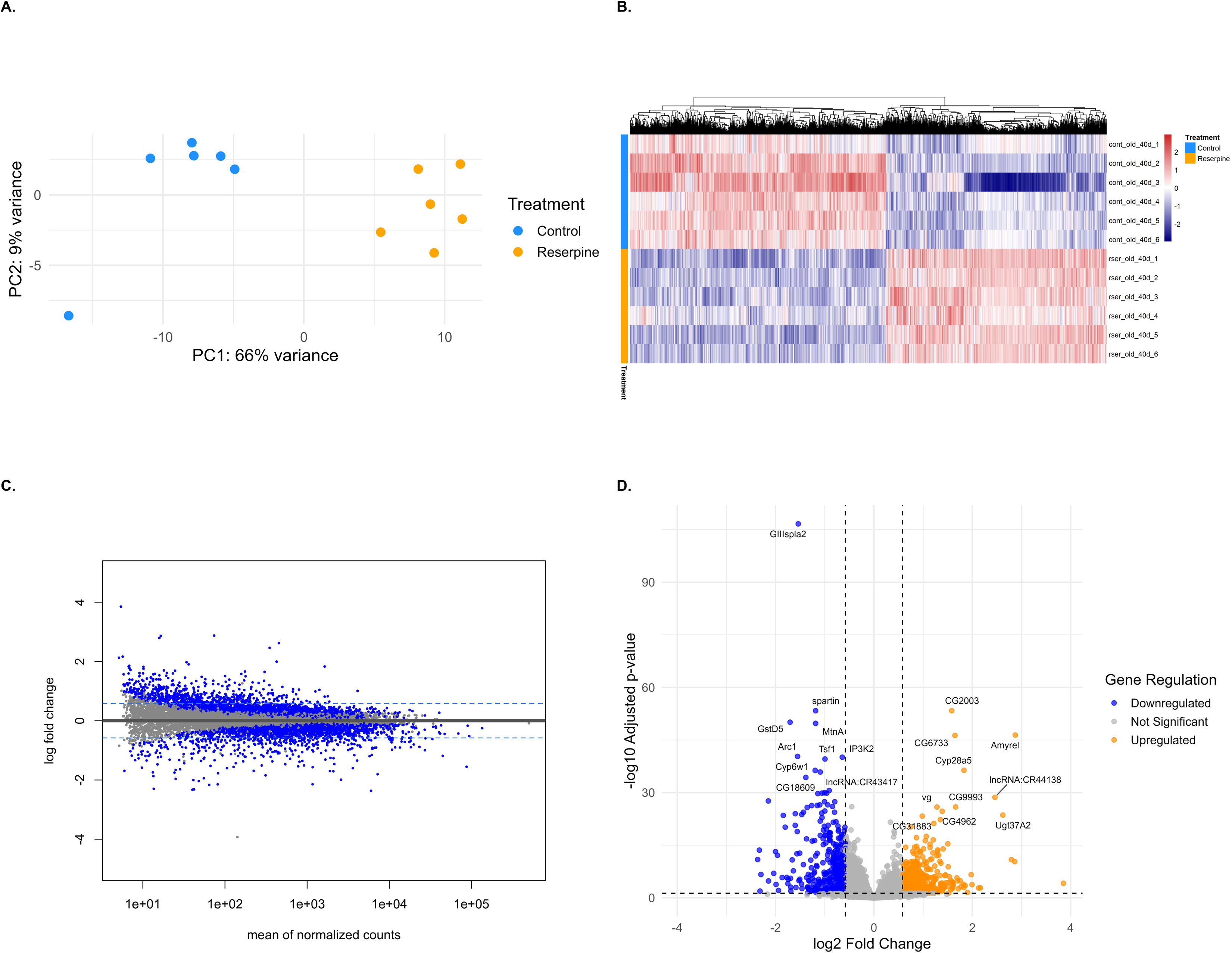
Transcriptomic profiling of aged flies (day 43) treated with reserpine compared to controls. (A) PCA plot showing transcriptome-wide differences between old control (red) and reserpine-treated old flies (blue). PC1 explains 66% of the total variance, showing a clear separation based on treatment. (B) Heatmap of DEGs (adjusted p-value < 0.05) with hierarchical clustering, displaying gene expression patterns across groups. (C) MA plot displaying log2 fold change versus mean normalized counts. Genes with |log2FC| > 0.58 and padj < 0.05 are marked in blue. Horizontal dashed lines represent ±0.58 thresholds. (D) Volcano plot of DEGs highlighting significantly upregulated (red) and downregulated (blue) genes, with the top 10 genes labeled by gene symbol and ranked by padj value.

### Reserpine Metabolic reprograming and stress-defense suppression in old flies

Over-Representation Analysis (ORA) of Gene Ontology (GO) terms including Biological Process (BP), Molecular Function (MF), and Cellular Component (CC) and KEGG pathways (**Figure 5A–D**), along with Gene Set Enrichment Analysis (GSEA) of the same categories **(Figure S3A-D**), revealed a strong downregulation of gene sets associated with fatty acid metabolism, inflammation, catabolism, proteostasis, peroxisomal and mitochondrial function and cytochrome P450-mediated detoxification.

**Figure 5.**
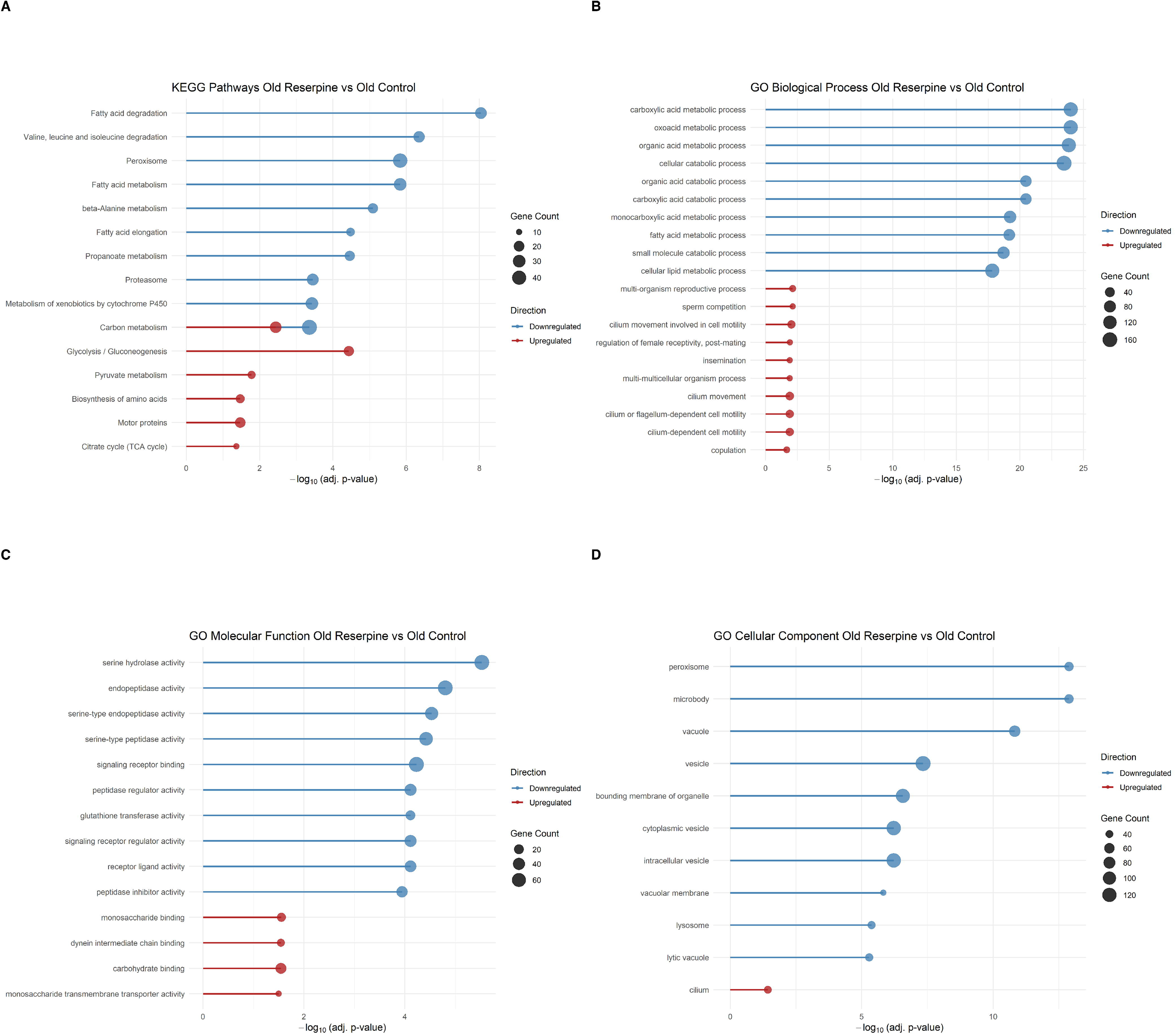
Over-representation analysis (ORA) of transcriptomic changes in reserpine-treated aged flies (Reserpine Old vs. Control Old). (A) KEGG pathway enrichment; (B) GO Biological Process; (C) GO Molecular Function; (D) GO Cellular Component. Downregulated pathways include fatty acid degradation, proteasome, peroxisome, and detoxification-related terms, while upregulated pathways highlight glycolysis, TCA cycle, and cilium-related processes. Analyses were performed using clusterProfiler. Each Dot size indicates the number of genes enriched per term, and position on the x-axis represents –log₁₀ (adjusted p-value).

In parallel, reserpine treatment weakly upregulated pathways related to glycolysis, pyruvate metabolism and the TCA cycle (**Figure 5A**), highlighting a metabolic reprogramming of glycolytic and mitochondrial energy metabolism. Together, these enrichment patterns indicate a transcriptional state that reshapes core bioenergetic processes while mitigating age-related metabolic and proteolytic stress. As seen in KEGG enrichment, carbon metabolism is both highly downregulated with greater gene count and significance in treated vs control flies, while parts of carbon metabolism are also relatively weakly upregulated further indicating a complex carbon metabolism transcriptional shift or reprogramming (**Figure 5A-B, S3A-B)**. This pattern suggests a form of mitochondrial remodeling in aged flies, favoring metabolic flexibility and reduced redox stress. Such shifts have been observed in other models of extended lifespan, including dietary restriction (DR), Indy mutants that mimic DR, and PRC2-deficient flies ^41–44^. A notable finding was the strong enrichment of serine hydrolase-related activity and serine type endopeptidase activity among the top four downregulated Molecular Function (MF) GO terms (**Figure 5B-C, S3B-C).** Serine hydrolases include proteases with roles in immune activation, stress response, and proteostasis, such as Persephone, a key immune protease in flies^45,46^. Dysregulated serine metabolism has been implicated in chronic inflammation ^47^ and aging-related diseases. ^47^

Collectively, these findings suggest that chronic reserpine exposure in aged flies leads to a transcriptomic state marked by downregulation of stress response and metabolic genes, potentially reflecting a shift toward a low-energy expenditure phenotype with implications for aging and longevity as also indicated by reduced activity in climbing assay.

### Reserpine Reshapes the Heat Stress Transcriptome in Young Flies

Next we compared the trancritpome profiles of 13 days young flies following heat stressed between control and chronic reserpine treated (12 days standard temperature + 24hr treatment at 31 °C). PCA revealed strong separation between groups, with PC1 explaining 70% of the variance, highlighting the dominant influence of reserpine in shaping transcriptomic responses to heat stress (**Figure 6A**). Similarly to aged flies, reserpine induced a marked shift in gene expression under heat stress conditions in young flies. The heatmap of DEGs (padj < 0.05, n = 2871) shows distinct clustering between treated and control samples, indicating a consistent reserpine effect on the transcriptome (**Figure 6B**). MA plots show again predominant downregulation of significantly altered genes, supporting a suppression of energy-demanding processes under reserpine treatment (**Figure 6C**). Volcano plots illustrate significant downregulation of genes involved in digestion (*Jon99Fii*, *Jon65Aiii*), detoxification (*Cyp6a8*) and sensory perception (Obp56d) (**Figure 6D**). Additional downregulated genes include *epsilonTry* and *Lsp1beta*, associated with protease activity and storage protein function, suggesting suppression of metabolic and sensory pathways during heat stress. Upregulated genes included *Prps* and *Lectin-galC1*, linked to biosynthetic and immune responses, and fork, a transcription factor involved in stress and longevity. TpnC41C, a calcium-binding protein, was also upregulated, potentially reflecting muscle-related stress.

**Figure 6.**
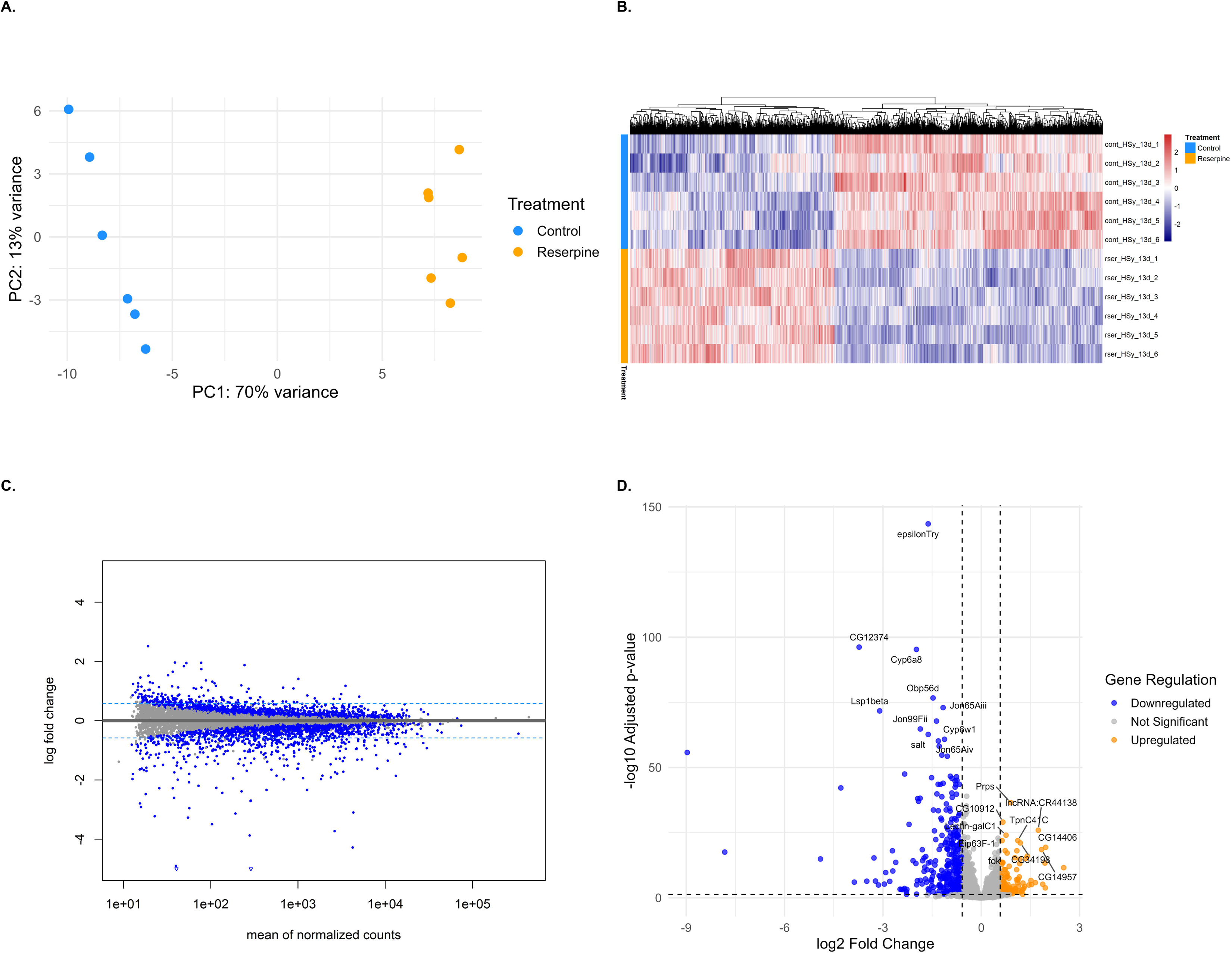
Transcriptomic profiling of young flies’ heat stress exposed flies. (A) Principal component analysis (PCA) showing distinct clustering of samples by treatment, indicating transcriptomic differences induced by reserpine under heat stress. (B) Heatmap of differentially expressed genes (DEGs; adjusted p-value < 0.05), illustrating patterns of gene expression across samples. (C) MA plot depicting the relationship between mean expression and log2 fold changes for all genes, with thresholds indicated for differential expression. (D) Volcano plot illustrating log2 fold change versus –log10 adjusted p-values of DEGs. Both significantly upregulated and downregulated genes are highlighted, with the top 10 genes labeled

### Reserpine impairs heat-stress response and suppresses stress-defense pathways

In young flies subjected to heat shock, reserpine treatment resulted in significant suppression of several key heat shock response genes. Notably, *Hsp23*, *Hsp26*, *Hsp70Bc*, *Hsp70Aa*, and *Hsp70Ab* were all significantly downregulated in reserpine-treated flies compared to heat-shocked controls (**Figure 7**). While *Hsp68* and *Hsp70Bb* were not significantly changed, they also showed a trend toward downregulation. These findings indicate that reserpine broadly impairs the transcriptional heat shock response, potentially compromising the fly’s capacity to manage proteotoxic stress. In flies, the maternal loading in embryos of Hsp23 mRNAs increases thermotolerance ^48^ and loss of *Hsp70* Genes including *Hsp70Ab* reduces thermotolerance and survival ^49^. Hsp70 group of genes (including *Hsp70Ab, Hsp70Bc, Hsp70Aa*) are among the most highly inducible and protective heat shock proteins in Drosophila, essential for survival under stress.^49^. *Hsp83* (the Drosophila homolog of Hsp90) is involved in protein folding and stabilization of key signaling proteins^50^. Small Hsps (*Hsp23, Hsp26, Hsp27*) are rapidly induced by heat and oxidative stress and help protect cells from damage^51^.Overall, several key pathways include those involved in proteasome, fatty acid metabolism, glutathione, and cytochrome P450 detoxification, (**Figure 8A-B, S4A-B) Peroxisome, lysosome,** serine hydrolase and peptidase activity (**Figure 8C-D, S4C-D)** were downregulated in Reserpine vs. control in young HS. Interestingly, these pathways, also suppressed in old treated flies, failed to upregulate in response to heat, compromising cellular defense and stress resilience as mentioned earlier. Simultaneously, there was a highly enriched upregulation of energetically costly processes such as ribosome and cytoplasmic translation, amino acid biosynthesis, synaptic signaling (**Figure 8A-B, S4A-B),** muscle contraction, ion channel activity, respiratory chain complex in drug treated HS flies (**Figure 8C-D, S4C-D).** This shift toward high-cost cellular activities may increase the vulnerability to heat stress, as energy is diverted from protective mechanisms to processes that further strain cellular resources.

**Figure 7.**
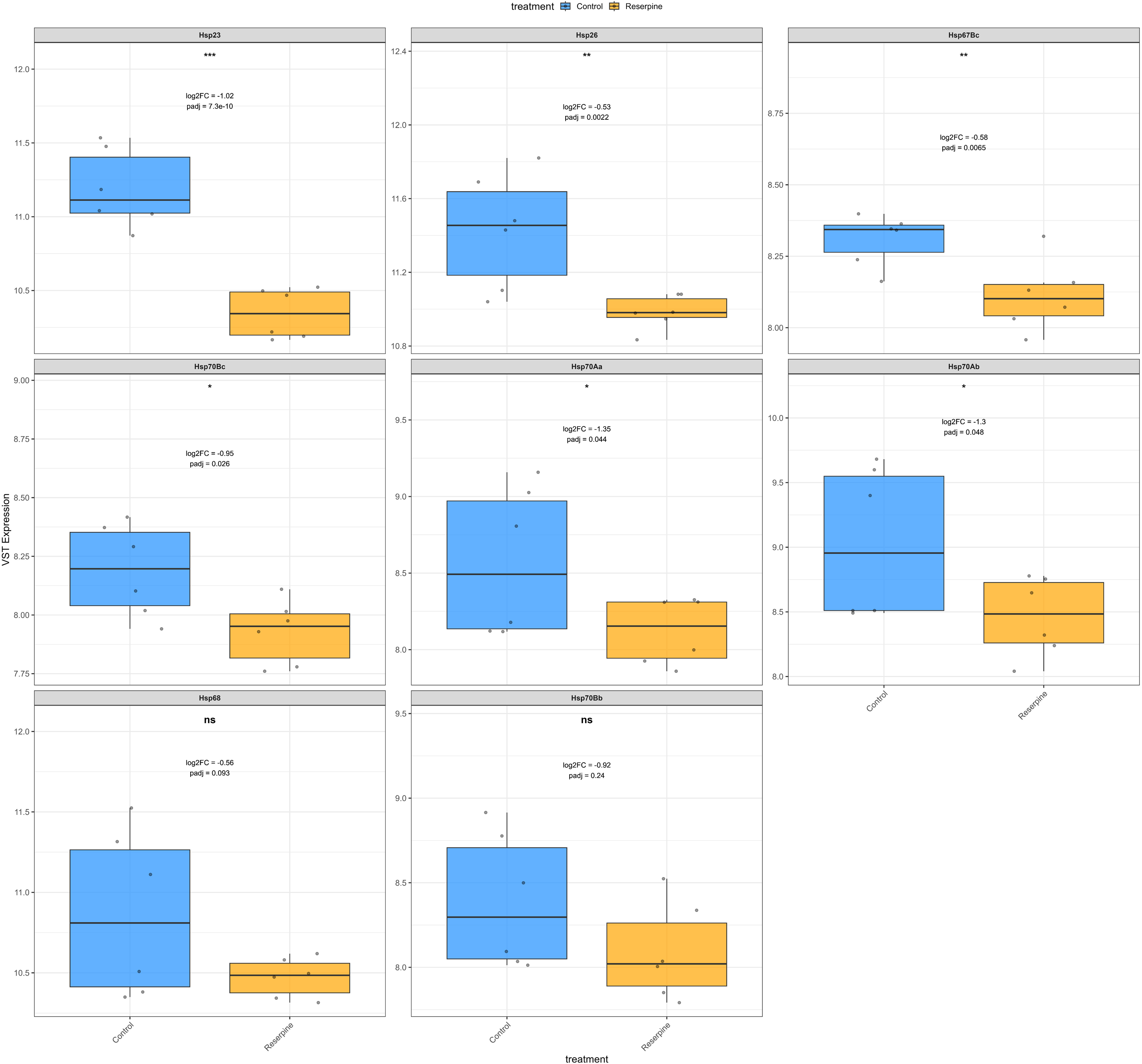
Expression of Heat Shock Genes in Young Flies After Heat Shock. Boxplots show variance-stabilized transformed (VST) mRNA expression levels of canonical Drosophila heat shock genes (young flies subjected to heat shock. Comparison is between control and reserpine-treated groups. Reserpine treatment leads to significant downregulation of heat shock gene expression. Annotations within each panel indicate the log2fold change, adjusted p-value, and difference in mean VST expression (ΔVST) between treatments. Only significant genes |log2FC| > 0.5 and padj < 0.05 are taken to be significant.

**Figure 8.**
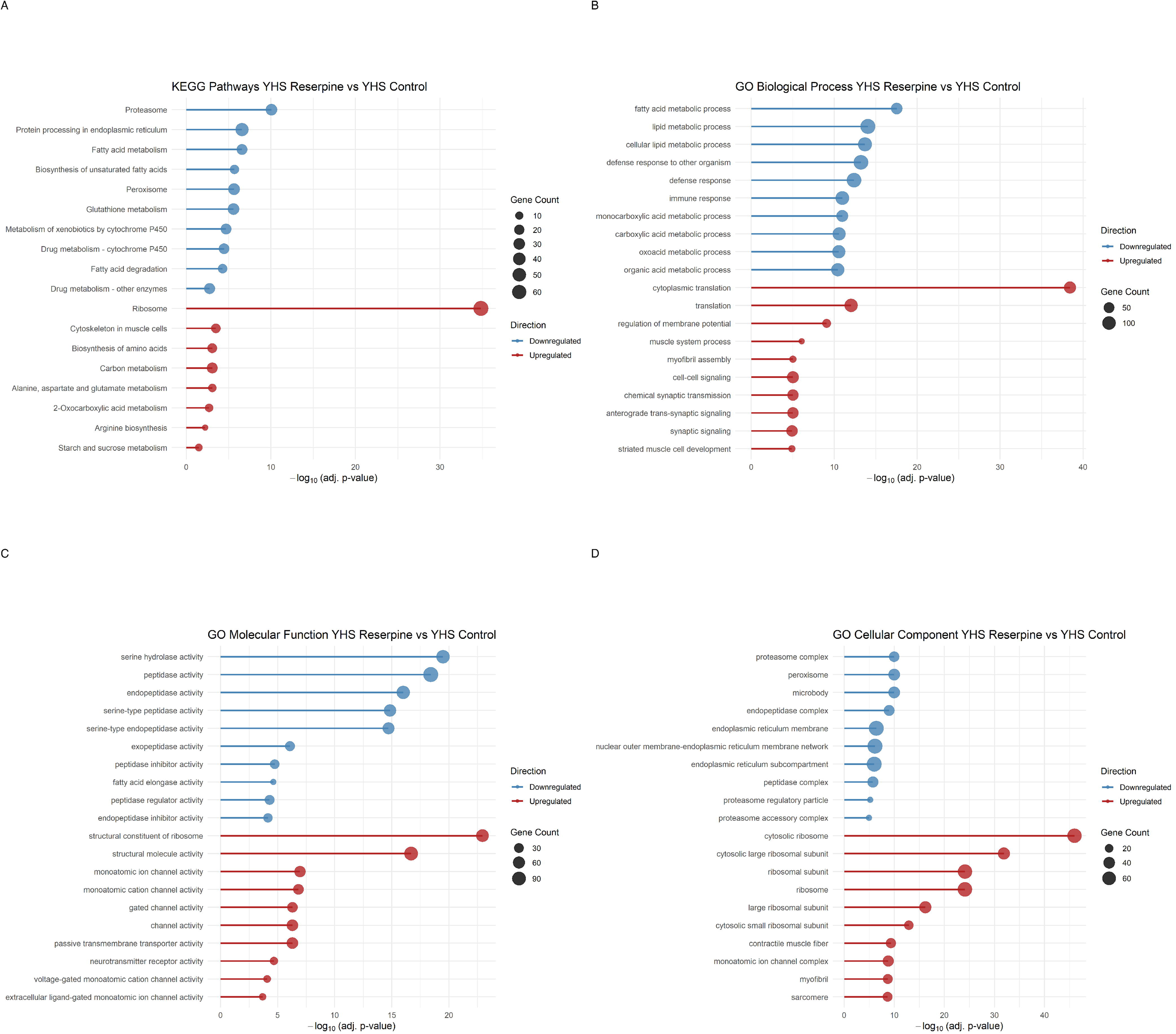
Over-representation analysis (ORA) of transcriptomic changes in reserpine-treated young flies under heat stress (YHS Reserpine vs. YHS Control). (A) KEGG pathway enrichment; (B) GO Biological Process; (C) GO Molecular Function; (D) GO Cellular Component. Downregulated terms include fatty acid metabolism, proteasome, glutathione activity, and immune/metabolic processes, while upregulated terms include translation, biosynthesis, ion channel activity, and sarcomere-related components. Analyses were performed using clusterProfiler. Dot size indicates gene count, and position on the x-axis represents –log₁₀(adjusted p-value). Color indicates direction: blue = downregulated, red = upregulated.

### Common features in treated flies among both cohorts

There are several common features in old and young flies under reserpine treatment such as metabolic suppression in old and maladaptive upregulation due to heat stress of processes that leads to lethality in heat shocked drug treated flies. Interesingly, we noticed that overlapping pathways were impacted by reserpine both in aged and in young flies heat stress when treated with the drug. Venn diagram revealed that 1427 genes (padj <0.05) were diffentially expressed in both the old and heat stressed groups. Of those, 844 genes were commonly down regulated by reserpine treatment while 447 genes were upregulated in both groups. Notably, 136 DEGs were differetially regulated upon treamtnet (old and heat stressed, but in opposite direction of regulation (**Figure 9A-D**). Enrichment of the common upregulated and downregulated genes revealed common terms similar with overall enrichments (**Figure S5**). Enrichment analyses thus confirm a broad suppression of protective, aging-linked pathways similar to observations in on old flies where terms like fatty acid, serine hydrolase and serine peptidase and glutathione metabolism, and metabolism via cytochrome P450 are downregulated in short treatment duration of 13 days and also in long term treatment of 43 days.

**Figure 9.**
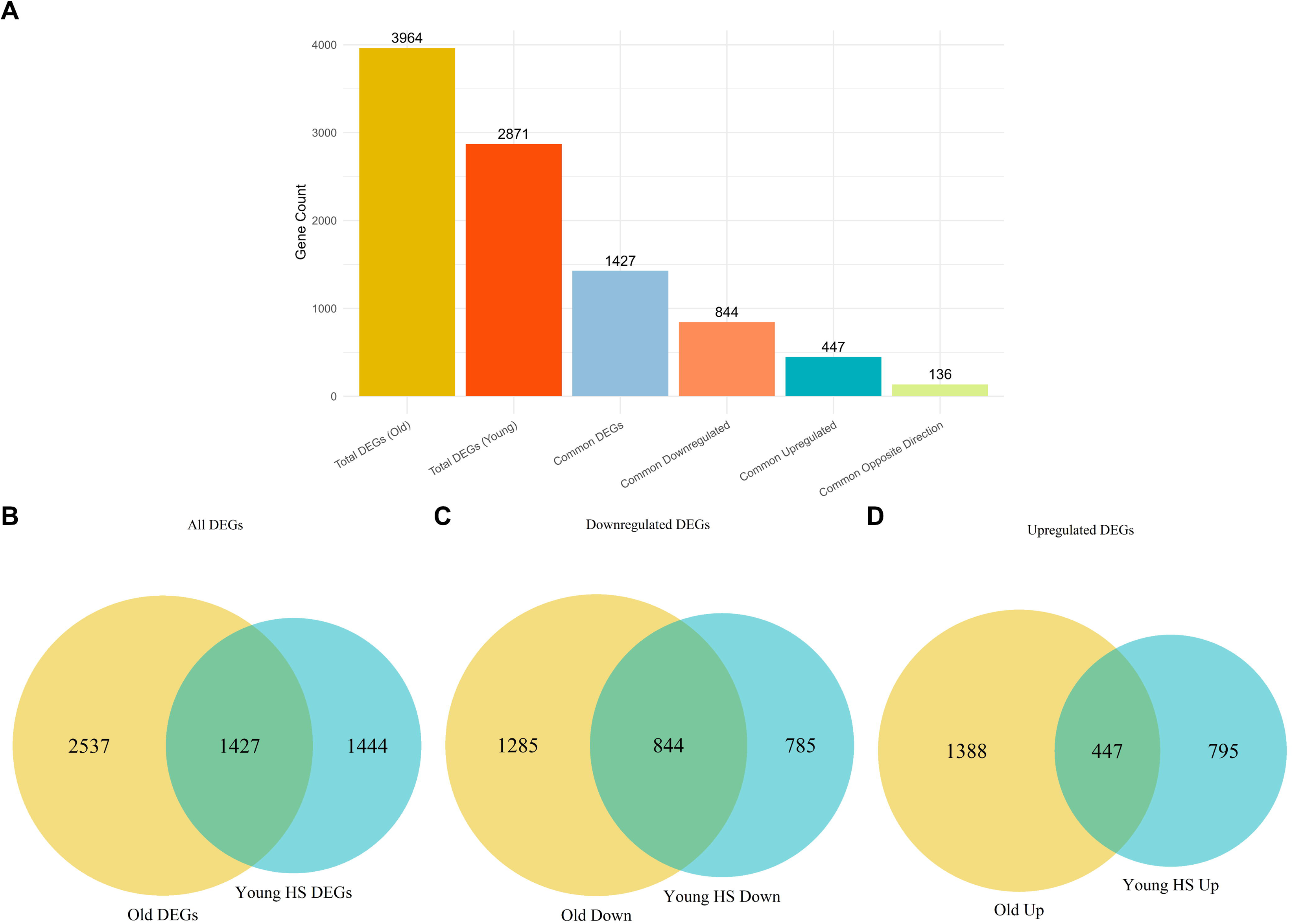
Shared transcriptional signatures between reserpine-treated aged flies and young flies under heat stress. (A) Bar plots showing common DEGs overall, up and down in both cases, and opposite in both cases. (B) Venn diagram showing overlap of all DEGs (padj < 0.05) between Reserpine Old vs. Control Old and Young Heat Stress Reserpine vs. Young Heat Stress Control. (C) Shared genes are separated into commonly downregulated subsets. (D) Shared genes are separated into commonly upregulated subsets

### Downregulation of known longevity genes in old flies

Lastly, we compared our differentially expressed genes (DEGs) with the GenAge Model Organism (https://genomics.senescence.info/genes/index.html) dataset for *Drosophila melanogaster*, which compiles 2,205 genes associated with aging or longevity based on genetic modification experiments reported in the literature. This curated set includes only genes that have a significant impact on lifespan and/or aging when genetically altered, while genes that reduce lifespan solely by inducing specific diseases without evidence of accelerated or premature aging are generally excluded to maintain a focus on the aging process. Within this dataset, genes are classified as either “anti-longevity” (n = 1,101), which promote aging or reduce lifespan, or “pro-longevity” (n = 545), which delay aging or extend lifespan ^52^. For *Drosophila melanogaster*, a total of 241 longevity genes with their function, longevity effect and study references were downloaded from their GenAge database website (https://genomics.senescence.info/genes/search.php?organism=Drosophila+melanogaster). We identified a consistent set of longevity-associated genes expression was significantly altered following reserpine treatment in aged Drosophila. To ensure both statistical and biological relevance, we filtered for genes that met the significance threshold (adjusted p-value < 0.05 and |logFC|>0.5). Only GenAge-annotated genes passing both thresholds were retained for downstream visualization and interpretation (**Figure 10**). A transcriptional suppression was observed in the old cohort, where reserpine treatment reduced expression of the nine pro-longevity genes, along with one anti-longevity gene. These annotations are based of anti and pro is based on 1-2 studies in the GenAGE database so does not provide a full picture and many of these gene can change depending on context of the treatment (**Figure 10**).

**Figure 10.**
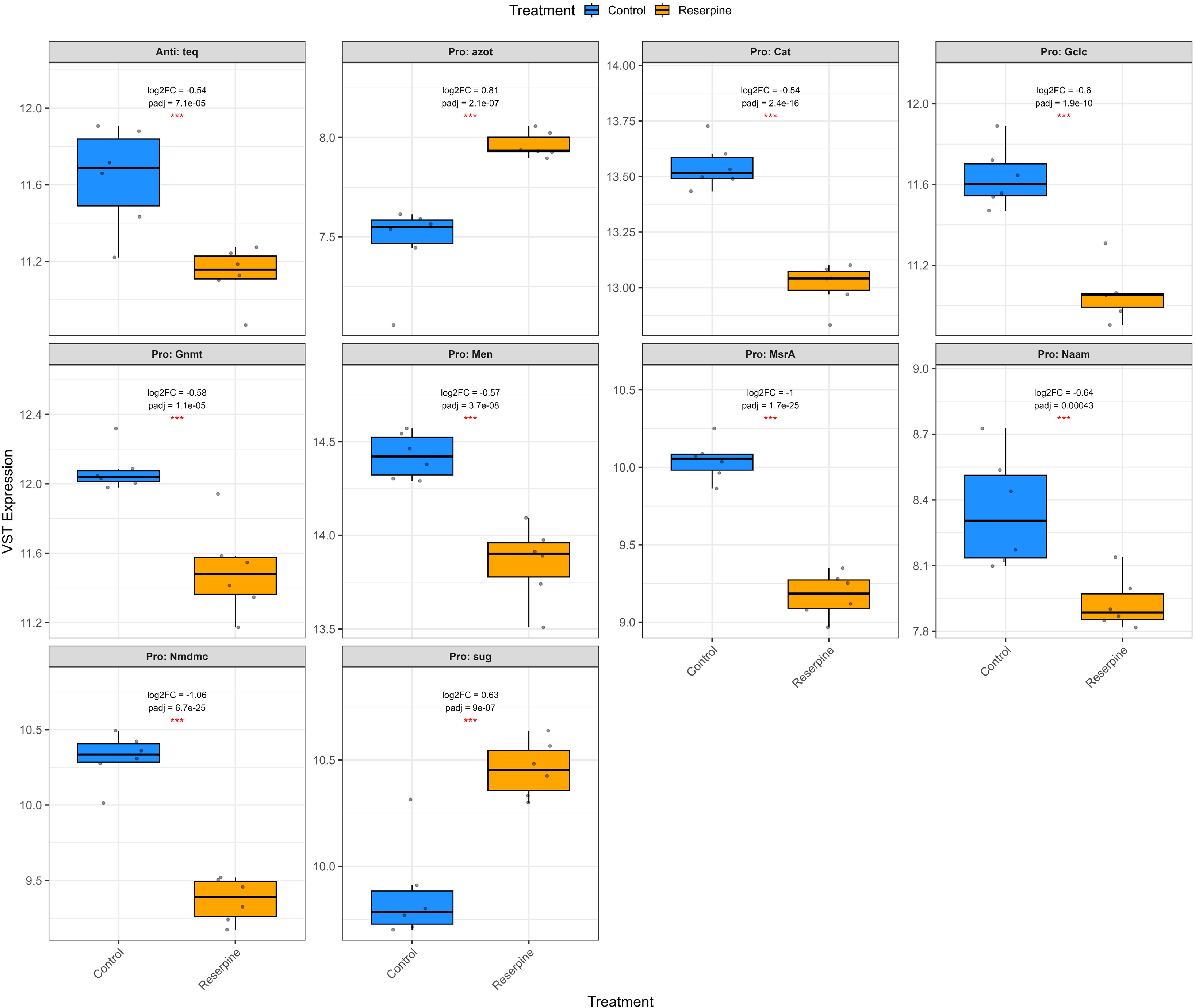
Differential expression of GenAge longevity-associated genes in old flies. Bar plots and clustered heatmaps depict significantly regulated GenAge genes (padj < 0.05, |VST difference| > 0.5). Many pro-longevity genes are downregulated in reserpine-treated flies, particularly in aged samples. Data represent normalized expression values.

## Discussion

Our study demonstrates that chronic dietary treatment with reserpine extends male drosophila lifespan while simultaneously impairing locomotor activity, reducing survival under heat stress, and suppressing metabolism and immune response including heat-shock response genes. These results highlight a classic geroscience trade-off: pharmacological interventions that enhance longevity may compromise stress resilience ^24^ and physical vigor, particularly under challenging conditions.

Previous studies have shown that depletion of serotonin or octopamine can stimulate a “low food” signal, activating dietary restriction (DR) - like pathway responses and also modulating metabolism and stress tolerance via mechanisms such as neuron–gut signaling and p38-MAPK ^53–55^. While mild depletion of either neurotransmitter may promote longevity by inducing metabolic restraint ^53,55^ excessive loss impairs stress resilience including heat, hypoxia, and pathogen defense due to suppressed stress responses^55–57^. In our study, strongly downregulated metabolic and stress-response pathways including reduced heat stress resistance in reserpine-treated flies are consistent with such monoamine neurotransmitter depletion–like states but via pharmacological VMAT inhibition, reflecting the trade-off between lifespan extension and stress vulnerability in treated flies. Additionally, depletion of dopamine in genetic DA-loss models shows impaired activity ^58^ which mirrors reduced locomotion and motor neuron activity VMAT mutants ^55,59^ reflecting in our findings with reduced climbing activity.

Our study identifies clear tradeoffs of reserpine-induced longevity, rendering the flies less active and displaying impaired heat stress resilience. Reserpine-treated flies failed to mount a canonical heat shock response, with significant suppression of inducible heat shock proteins (HSPs) such as Hsp23, Hsp26, Hsp67Bc and others. HSPs are critical for thermotolerance, protein refolding, and cellular survival during proteotoxic stress and aging ^40,48,49^. The suppression of detoxification pathways to activate essential adaptive responses contributes to increased mortality during heat exposure. As observed in an earlier study by Bressan, et al. ^37^, reserpine treatment significantly reduces heat stress tolerance in Drosophila melanogaster, impairing their ability to avoid noxious heat and decreasing locomotor activity, likely through disruption of monoaminergic signaling pathways. This pharmacological intervention compromises the flies’ stress resilience, making them more susceptible to heat-induced damage ^37^. Furthermore, in our study, immune, and inflammatory signaling components, including those related to innate immunity and lysosomal proteases, were suppressed. This aligns with the concept of inflammaging, whereby chronic low-grade immune activation contributes to tissue decline ^60^.

Reserpine induced a pronounced transcriptomic shift in old as well as young HS cohorts, marked by broad suppression of metabolic, immune, and stress-response genes. Mechanistically, this suppressed metabolic state seems to be analogous to the suppression of mTOR i.e., master regulator of cellular metabolism. Metabolic suppression via mTOR depletion is shown to promote longevity ^61^, but in our study it is achieved via neuromodulation through reserpine treatment. Consequently, with metabolic suppression, few pro-longevity genes were also downregulated in treated flies ^52^ (**Figure 10**). At first glance, this might seem counterintuitive but it is known that pro-longevity genes can be related to stress-resistance or anti-oxidant genes^62^. Anti-oxidant gene suppression might reflect a reduced demand for oxidative stress defense due to broad reduction in metabolic load, similar to mTOR suppression or sustained caloric restriction ^61,63,64^. This can be observed as downregulated carbonic and fatty acids metabolism enrichment (**Figure 5A, B**). Reduced energy demand and/or need is also reflected in reduced activity of flies in climbing assays (**Figure 2**).

Interestingly, two key pathways that were enriched in downregulated terms by reserpine treatment in both old and young HS group were serine metabolism (hydrolase and peptidase) and glutathione transferase (GST) activity. Serine supplementation in *C.elegans* and yeast has been shown to increase lifespan, while in pre-clinical rodent studies show reductions in both circulating and hippocampal serine levels with age ^47^. Aging in Drosophila is linked to shift from glycolysis to serine metabolism and purine metabolism. In contrast, long-lived models such as PRC2 mutants maintain youthful serine metabolism and avoid such metabolic shifts ^39,44^, suggesting that reducing serine peptidase or hydrolase activity may be longevity-promoting by allowing higher serine availability by suppressing its metabolism. Similarly suppressed glutathione transferase (GST) activity which is central to oxidative stress defense ^65^, indicates immune suppression which could indicate less oxidative burden or alternative antioxidant mechanisms in play. Reserpine’s ability to suppress this immune-metabolic axis may be central to its pro-longevity effect. Collectively, these point to a suppression of energy-intensive and stress-inducing catabolic processes, often upregulated during aging and chronic immune activation ^1,66^.

In contrast to the widespread metabolic suppression, key glycolytic and mitochondrial metabolic pathways were mildly upregulated, including Glycolysis/Gluconeogenesis, TCA cycle, and Amino acid biosynthesis. These shifts may reflect a reorganization of mitochondrial energy metabolism toward substrates such as carbohydrates and select amino acids, with reduced reliance on fatty acid β-oxidation or branched-chain amino acid catabolism both of which generate greater oxidative stress ^67^. Many common suppressions in both young HS and old group is shown in **Figure S5**, like Peroxisome, Glutathione metabolism, Drug and Xenobiotic metabolism by Cytochrome P50, Antioxidant activity. In old flies, a major but weakly upregulated category enriched across GO CC and GO BP is cilium, and cilium-dependent cell motility, insemination, and courtship behavior in treatment group. These enrichments hint at reserpine preserving or reactivating neurological or sensory-motor capacity in aged flies. Given that cilia are central to chemosensation and behavioral circuits in Drosophila ^68,69^, this may reflect a compensatory mechanism for maintenance of behavior vigor in context of chemosensation and response to stimuli, not simply reproductive investment due to reduced activity in treated flies.

While this study focused on flies, similar observations with reserpine have been made in worms. Notably, Saharia, et al. ^3^ showed that reserpine extends lifespan in C. elegans via mechanisms involving the dopamine receptor dop-3 and the RNAi pathway regulator eri-1 ^3^. However, our findings indicate that in flies, reserpine acts primarily through suppression of metabolic and inflammatory stress, coupled with energy reprogramming. This may highlight species-specific divergence in the molecular architecture of lifespan regulation.

From the list of gene compared (**Table S1**), we found that in both old and young heat-shocked (YHS) flies, *Act88F*, a gene encoding an indirect flight muscle actin ^70^, was significantly upregulated in reserpine-treated flies indicating a consistent transcriptional response related to muscle remodeling ^71^. Another gene, *TpnC41C* (Troponin C at 41C), encodes a muscle-specific protein primarily expressed in the jump muscles ^72^ is upregulated in both young HS and old flies. Both genes’ upregulation could be a compensatory mechanism to reduced locomotor ability. Moreover, *mt: ND3* (mitochondrial NADH-ubiquinone oxidoreductase chain 3) which is subunit of complex I of the mitochondrial respiratory chain involved in energy metabolism^73^, was also significantly downregulated, potentially reflecting age-related susceptibility of mitochondrial function to monoaminergic disruption in old flies. Suppression of complex 1 of the Electron transport chain is associated with longevity in oocytes by avoiding ROS generation^74^. Genes like *ple* which encodes a tyrosine hydroxylase^75^, the first and rate-limiting step in the synthesis of dopamine ^76^ and *TyrR* when overexpressed has neuromodulatory effect of male courtship behavior leading to high mating behavior ^77^.

Taken together, the transcriptomic profiles demonstrate that reserpine triggers a marked, condition-specific suppression of metabolic, immune, and stress-response pathways in both aged and heat-stressed flies. This effect is reflected in the consistent downregulation of key genes and the clear separation of transcriptomic signatures between treated and control groups, thus supporting the conclusion that reserpine promotes a immune suppressed and low energy expenditure state in flies under both physiological aging and acute stress conditions. In summary, reserpine’s effects in Drosophila illuminate the evolutionary trade-offs that may come with current longevity interventions. By isolating beneficial mechanisms and refining pharmacological approaches, it may be possible to promote healthy lifespan extension while minimizing adverse outcomes.

### Limitations of the study

Despite the strengths of our integrative approach combining lifespan assay, climbing activity assay, heat stress tolerance assay, transcriptomic, and pathway analyses our study has several limitations. Our focus was on male flies only. We also did not determine reproductive output in females, which are often affected by monoamine depletion and may contribute to the observed trade-offs. Finally, while our RNA-seq provides a snapshot of global gene expression, complementary proteomic and metabolomic data would provide a more complete picture of the molecular consequences of reserpine treatment.

### Future Directions

Our findings underscore both the benefits and complexity of repurposing reserpine like monoaminergic signaling altering drugs as a geroprotective agent. As an FDA-approved VMAT inhibitor, reserpine modulates monoaminergic signaling and may mimic some features of dietary restriction or stress hormesis mechanisms linked to lifespan extension. However, the observed trade-offs, include reduced stress resilience, reduced locomotor activity. This duality highlights the necessity for careful monitoring when targeting similar pathways for aging interventions. Mechanistically, future studies should dissect which monoaminergic circuits dopaminergic, serotonergic, or octopaminergic are most critical for mediating reserpine’s effects specific to Fruit flies. Targeted manipulation of these pathways may enable the development of interventions that maximize longevity benefits while minimizing adverse effects. To achieve this, comprehensive phenotyping including behavioral, metabolic, and reproductive assays should be integrated with transcriptomic and proteomic analyses to elucidate the systemic consequences of chronic monoamine depletion. Cross-species validation, particularly in mammalian models, will be essential to assess the translational potential and safety of reserpine related compounds. Given the context-dependent nature of reserpine’s effects, systematic investigation of dose-response relationships, intervention timing will be crucial. These parameters influence the balance between lifespan extension and health span, as observed with other agents such as anti-diabetic and anti-obesity drugs like metformin or semaglutide-based drugs (Ozempic).

## Materials and Methods

*Drosophila melanogaster* (Canton-S wild type) were reared at 25°C under a 12:12 hour light: dark cycle. For all experiments, males aged 1–2 days at the start were used. Reserpine (Sigma-Aldrich, Cat. No. 83580; MW 608.68 g/mol) was dissolved in dimethyl sulfoxide (DMSO) to generate a 100 mM stock solution (760 mg in 12.5 mL DMSO). After autoclaving and cooling the fly food to approximately 60°C, reserpine was added to achieve final concentrations of 0 (DMSO control), 200, 800, 1000, or 1500μM, as indicated for each experiment.

Fly food was prepared with a standardized cornmeal–soy–molasses medium (containing 3.21 L distilled water, 30 g soy flour, 260 g corn flour, 60 g active dry yeast, 26 g agar, 260 g molasses, and 130 g malt extract per batch), homogenized and autoclaved at 121°C for 20 minutes. After cooling to ∼60°C, 83 mL of 10% (w/v) methylparaben (in ethanol) and 59 mL acid mix (60 mL 85% phosphoric acid, 418 mL propionic acid, distilled water to 2 L) were added, and the medium was dispensed aseptically (8 mL per vial) and allowed to solidify.

### Survival Assays

For survival assays under standard conditions,1-day old flies were placed in three biological replicates with n ∼ 35 flies per vial and N ∼ 105 flies per condition were maintained at 25°C and 60% humidity on reserpine-supplemented media and flipped on every alternate day. Mortality was recorded three times a week. For heat stress survival experiments, 1-day old flies were placed in three biological replicates of 35 flies per group were transferred to reserpine-supplemented media and incubated at 31°C inside incubator, with flies flipped daily with dead flies recorded until all flies had perished.

### Climbing assay

To evaluate locomotor function, a negative geotaxis climbing assay was performed on adult Drosophila melanogaster following chronic drug exposure. Flies were maintained on food containing 0 µM (control), 1000 µM, or 1500 µM of the test compound for 12 consecutive days, and the climbing assay was conducted on day 13.

Each treatment group consisted of six biological replicates, with n ∼ 25–30 flies per replicate. For the assay, flies were gently transferred into clean, transparent plastic vials measuring 9 cm in height, pre-marked into five equal quadrants (each 1.8 cm high) based on ^35^. To initiate the assay, flies were tapped to the bottom three times to standardize the starting position and stimulated to climb. Each trial was video recorded for 30 seconds, and the climbing position of flies was scored at the 10-second mark after 3 taps. At that point, the number of flies in each of the five vertical quadrants was recorded.

A Weighted Climbing Index (WCI) was calculated for each replicate using the formula:

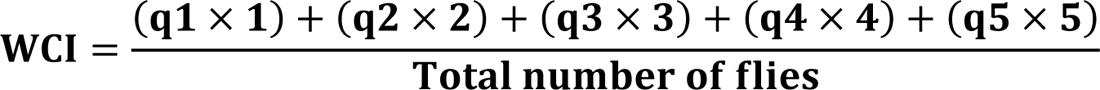

where qi is the number of flies present in the i^th^ quadrant from bottom (1) to top (5) of the climbing vial. A higher WCI indicates better climbing performance, as more flies reach higher quadrants.

Statistical analysis was performed in GraphPad Prism using one-way ANOVA followed by Tukey’s multiple comparisons test, comparing all treatment groups. Results are presented as mean ± SEM, with significance set at p < 0.05.

### RNA Extraction and Sequencing

Flies were collected at the following time points: old (43 days), and after young heat stress (12 days at 25°C followed by 24 hours at 31°C) in fly food with 1000µM Reserpine.

Flies were flash-frozen in liquid nitrogen, and total RNA was extracted using the Monarch Total RNA Miniprep Kit (NEB T2010S). The RNA concentration was analyzed using NanoDrop and the RNA integrity was assessed on the 2100 Bioanalyzer system using an Agilent RNA 6000 Nano Kit (Agilent Technologies). Strand-specific mRNA libraries were prepared from 1 μg total RNA using the Illumina Stranded mRNA Prep Ligation kit according to the manufacturer’s recommendations (Illumina). Library quality and fragment size were verified using Agilent DNA 1000 Kit (Agilent Technologies). Libraries were quantified using a Qubit dsDNA HS Assay Kit (Invitrogen), normalized to 10 nM, pooled, denatured to 750 pM, and paired-end sequenced (2×101bp) on the NextSeq 2000 system using P3 flowcell (Illumina) at the FBN sequencing facility, Dummerstorf, Germany.

### Data Analysis

Raw sequencing data (BCL files) were converted to FASTQ format using DRAGEN BCL Convert v3.10.11. Quality assessment was performed with FastQC v0.11.9. Adapter trimming, low-quality read filtering (mean Phred <20), and removal of reads <20 bp were performed using Trim Galore v0.6.10. Reads were aligned to the D. melanogaster BDGP6.46 reference genome (Ensembl r113) using HISAT2 v2.2.1. Gene-level counts were generated using HTSeq v2.0.2 (“union” mode), and genes with fewer than 10 total counts were excluded. Count matrices were imported into R via tximport and analyzed with DESeq2 v1.32.0. Data were normalized using the median-of-ratios method, and variance-stabilizing transformation was applied. Differential expression was tested using Wald tests with Benjamini–Hochberg multiple testing correction (padj <0.05). Differentially expressed genes (DEGs; padj < 0.05) were annotated and subjected to functional enrichment analyses using clusterProfiler v4.0, DOSE, enrichplot, and org.Dm.eg.db. Over-Representation Analysis (ORA) was performed for GO (Biological Process, Cellular Component, Molecular Function) and KEGG pathway terms using enrichGO() and enrichKEGG() functions (BH-adjusted p < 0.05; minGSSize = 10; maxGSSize = 500). Gene Set Enrichment Analysis (GSEA) was conducted using ranked log2 fold-changes.

Data visualization was carried out using ggplot2, including volcano plots and dot plots. Box plots were used to visualize expression differences for selected gene sets (GenAge, Heat Shock Genes) using VST expression and annotated with padj and |logFC| values. Genes were considered biologically relevant if they met both statistical (padj < 0.05) and expression (|logFC| ≥ 0.5) thresholds.

### Statistical Analysis

All statistical analyses were conducted using R version 4.4.3 and GraphPad Prism (10.5.0). Survival data were analyzed using Kaplan–Meier survival curves, with log-rank tests used for group comparisons in R. Cox proportional hazards models, stratified by replicate, were used to calculate hazard ratios (HR) and 95% confidence intervals (CI).

For RNA-seq data, differential gene expression was assessed using the Wald test, and statistical significance was determined using the Benjamini–Hochberg correction for multiple testing, with a false discovery rate threshold of padj < 0.05.

Climbing assay performance was evaluated using the Weighted Climbing Index (WCI). WCI values were analyzed in GraphPad Prism using one-way ANOVA, followed by Tukey’s multiple comparisons test to determine statistically significant differences between treatment groups

### Data Availability

Raw fastq files and metadata are available in the ArrayExpress database (http://www.ebi.ac.uk/arrayexpress) under accession number E-MTAB-15429 (https://www.ebi.ac.uk/biostudies/ArrayExpress/studies/E-MTAB-15429?key=255702e5-c660-407a-9cd9-cdb2ea27ed35).

## Supporting information

Aging cohort

Young heat shock cohort

## Acknowledgements

We thank Verena Hofer-Pretz for her invaluable technical assistance in maintaining fly cultures, preparing reserpine-containing media, and ensuring good laboratory conditions. We are also grateful to Leon Borowski at the FBN Next Generation Sequencing (NGS) Facility for library preparation for the RNA-seq experiments. We acknowledge Anuroop Venkatasubramani, Surya Hembrom, and Akshay Patil for their guidance on transcriptomic data analysis. Additionally, we acknowledge the use of AI tools (ChatGPT-4o, and Perplexity AI) which were used in refining R code. All AI-generated code and sources were carefully verified and reviewed.

## Supplementary figures

**Figure S1.** Pilot study assessing median survival at lower reserpine concentrations. Kaplan–Meier survival curves for male flies chronically treated with (0, 25, 50, 100, 200, 400, and 800 µM). Median survival values were: 49 (0 µM), 49 (25 µM), 51 (50 µM), 49 (100 µM), 53 (200 µM), 49 (400 µM), and 53 (800 µM). Each treatment group included 3 independent biological replicates each containing an average of n ∼ 35 flies per vial. Overall group differences were significant (log-rank test, p < 0.0017).

**Figure S2.** Follow-up pilot study showing survival at selected concentrations and a late-life intervention group. Kaplan–Meier survival curves for male flies chronically treated with Median survival values were: 51 (0 µM), 51 (200 µM), 51 (800 µM), 55 (1000 µM), and 39 (0-800 µM; reserpine introduced at day 31). These data suggest that timing of exposure when young is critical for the beneficial effects of reserpine. Each treatment group included 3 independent biological replicates each containing an average of n ∼ 35 flies per vial. Overall group differences were significant (log-rank test, p < 0.0001).

**Figure S3.** Gene Set Enrichment Analysis in Old Confirm global downregulation of immune and stress response pathways. Whole gene set is used in GSEA pAdjustMethod = “BH”, pvalueCutoff = 0.05, minGSSize = 10, maxGSSize = 500,

**Figure S4.** Gene Set Enrichment Analysis in Young Confirm global downregulation of immune and stress response pathways from early age. Whole gene set is used in GSEA pAdjustMethod = “BH”, pvalueCutoff = 0.05, minGSSize = 10, maxGSSize = 500,

**Figure S5.** Comparison of transcriptomic changes between aging and heat shock reserpine vs control flies. Enrichment of Overlapping downregulated DEGs (pvalueCutoff = 0.05, minGSSize =10, maxGSSize = 500) Enrichment of Overlapping Upregulated DEGs (pvalueCutoff = 0.05, minGSSize = 10, maxGSSize = 500)

**Table S1:** List of gene tested, related to monoamine depletion due to Reserpine Treatment.

