## Supplementary figures and images for "Reserpine prolongs lifespan but compromises heat-stress resilience in *Drosophila melanogaster*"

### Figure S1

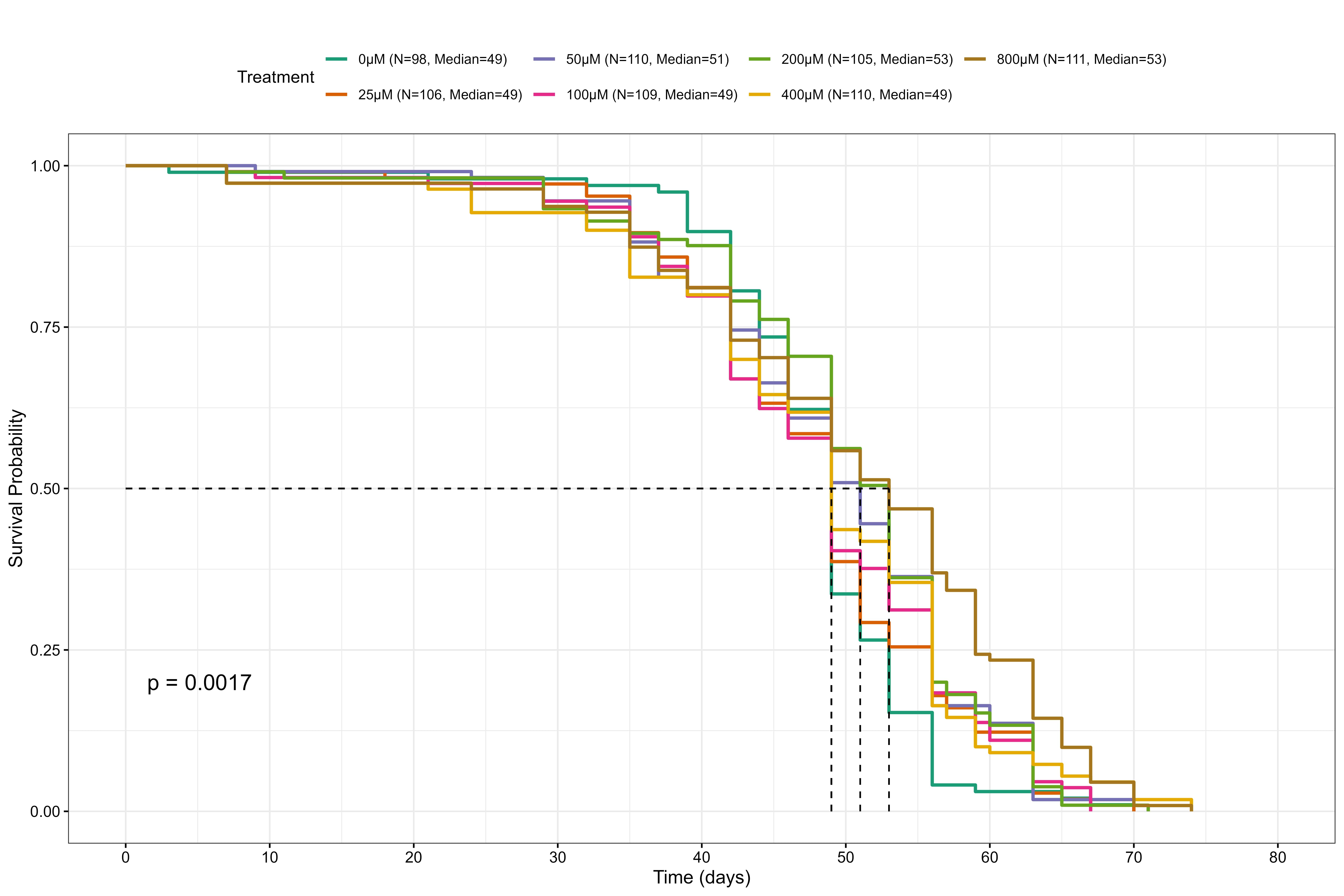

### Figure S2

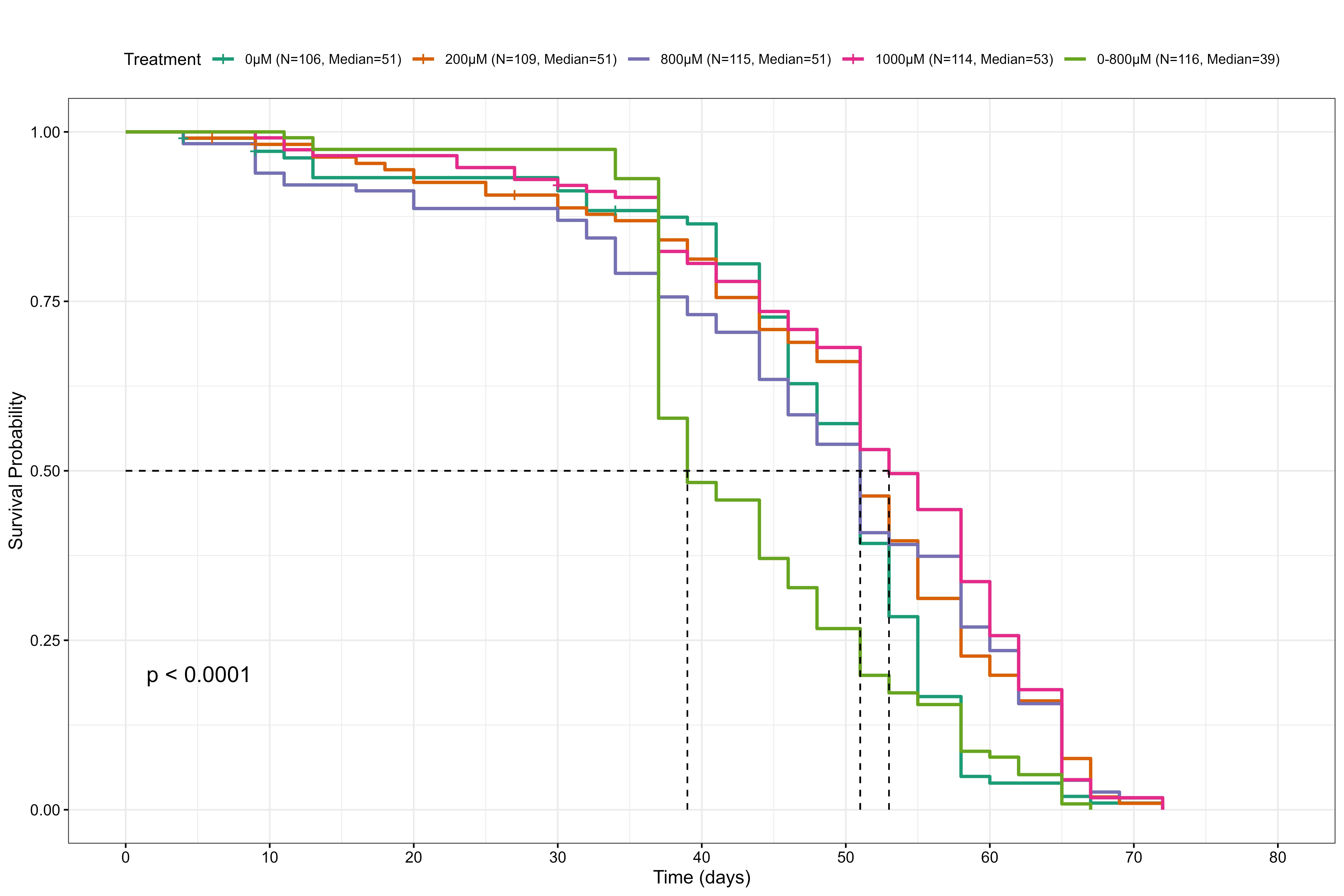

### Figure S3

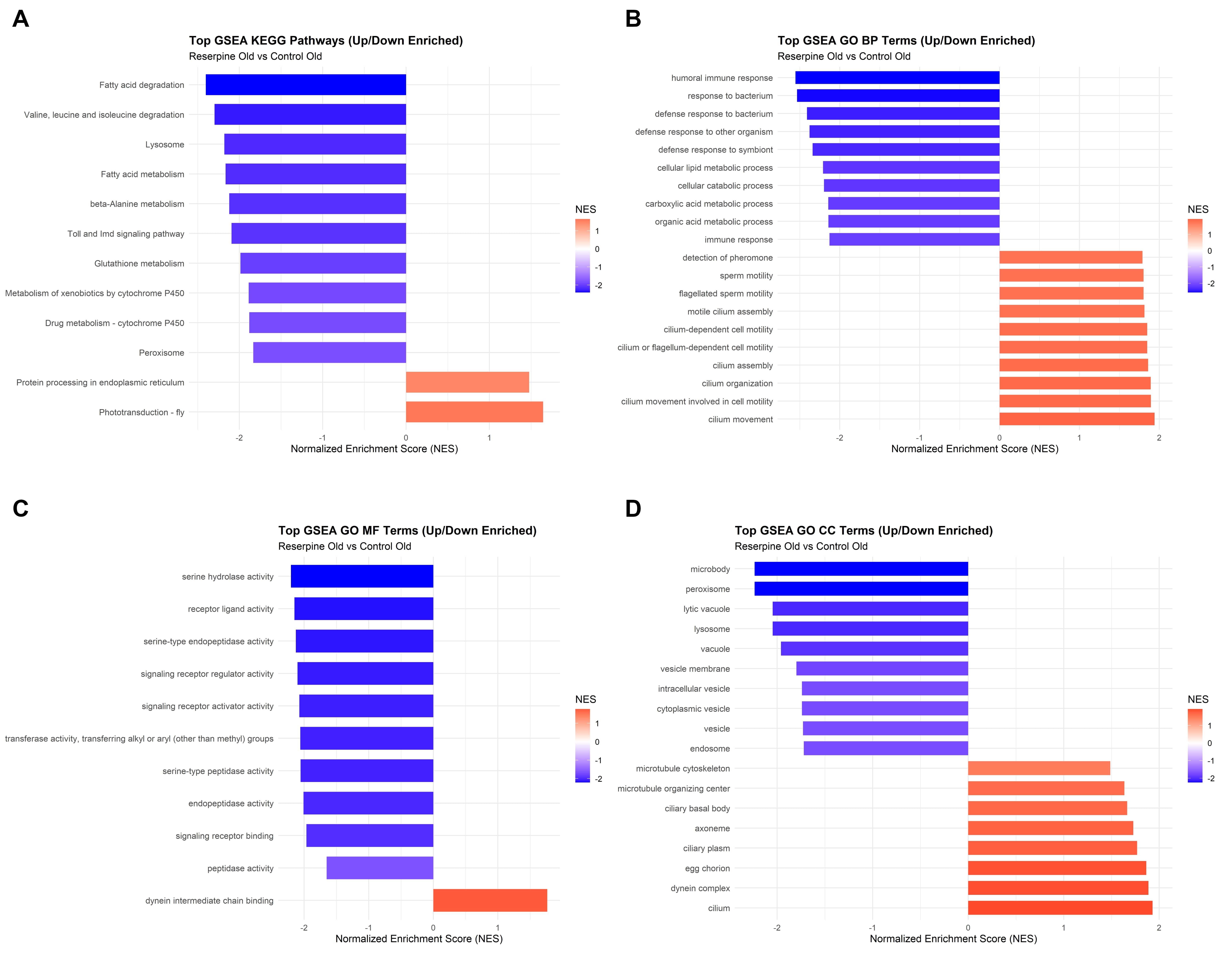

### Figure S4

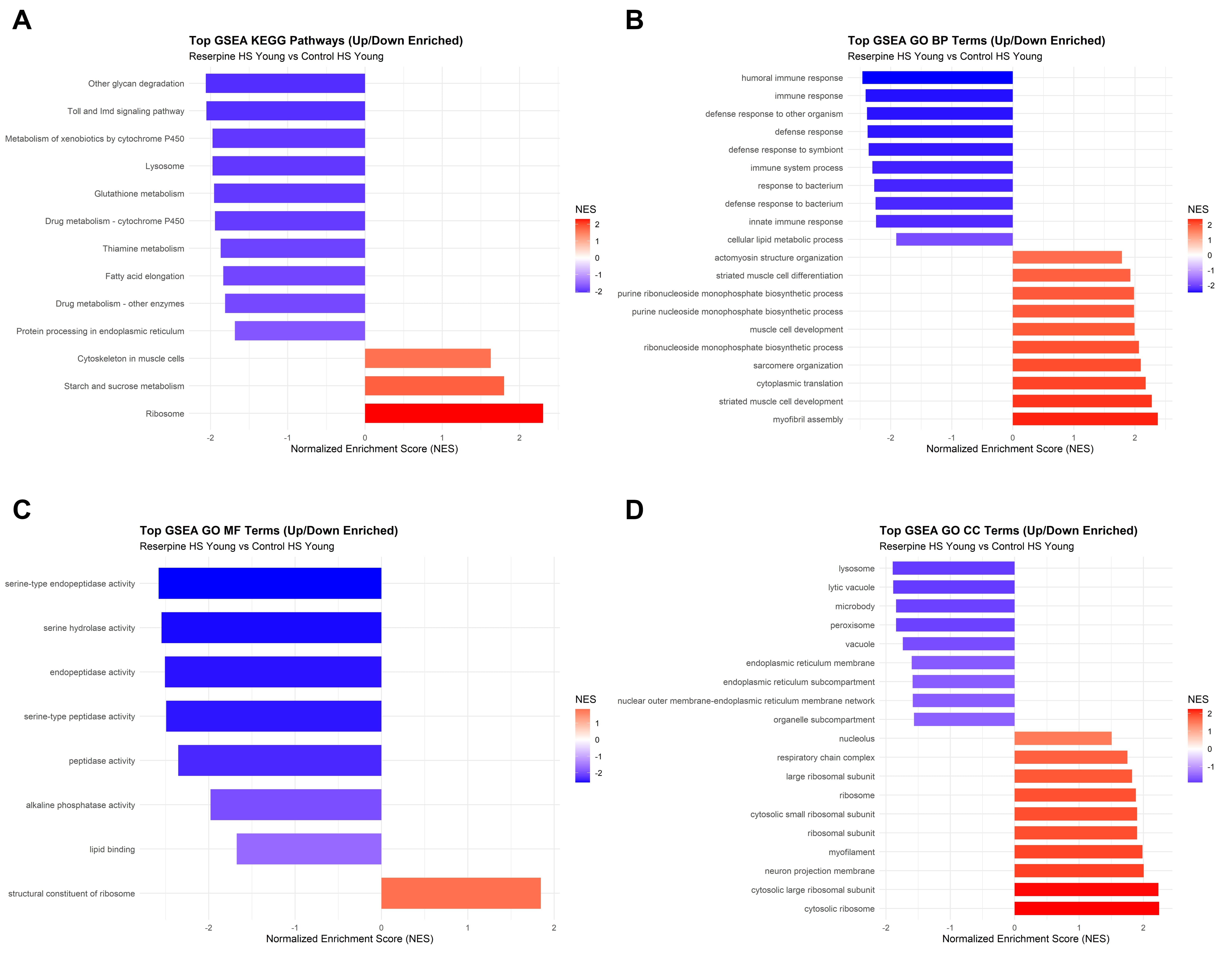

### Figure S5

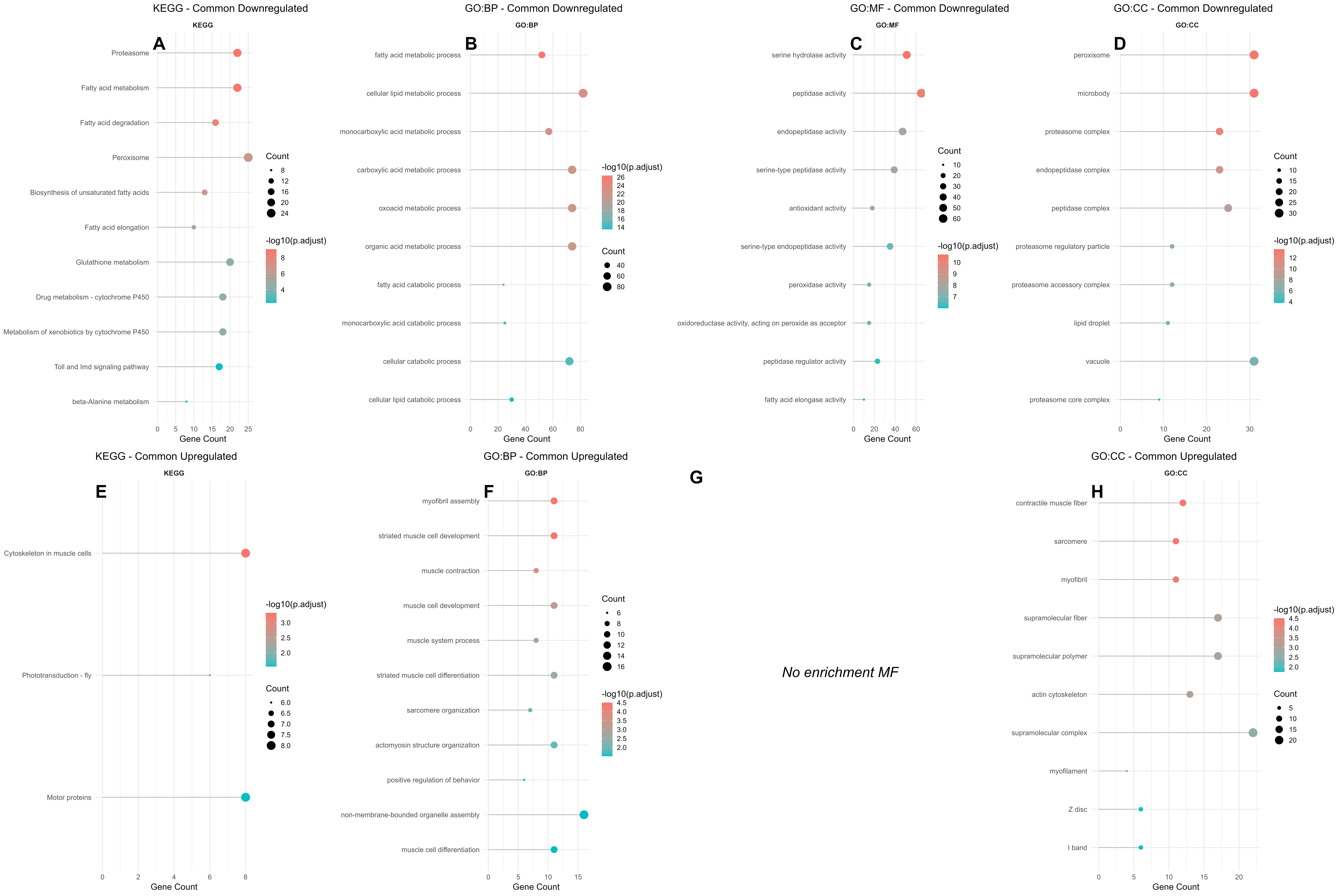
