## Supplementary material for "Reserpine prolongs lifespan but compromises heat-stress resilience in *Drosophila melanogaster*": Table S1

| Gene Symbol | FlyBase Gene ID | Functional Category |
| --- | --- | --- |
| ple | FBgn0005626 | Tyrosine hydroxylase (dopamine synthesis) |
| Punch | FBgn0003162 | GTP cyclohydrolase I (cofactor biosynthesis) |
| Ddc | FBgn0000422 | Dopa decarboxylase (dopamine/serotonin synthesis) |
| Tph | FBgn0262139 | Tryptophan hydroxylase (serotonin synthesis) |
| Tdc2 | FBgn0050446 | Tyrosine decarboxylase (tyramine/octopamine synthesis) |
| Vmat | FBgn0260964 | Vesicular monoamine transporter |
| DAT | FBgn0034136 | Dopamine transporter |
| SerT | FBgn0010414 | Serotonin transporter |
| Ebony | FBgn0000527 | Dopamine metabolism (β-alanyl-dopamine synthesis) |
| CG10433 | FBgn0034638 | Predicted monoamine transporter |
| Dop2R | FBgn0053517 | Dopamine receptor 2 |
| 5-HT1A | FBgn0004168 | Serotonin receptor 1A |
| 5-HT1B | FBgn0263116 | Serotonin receptor 1B |
| 5-HT2A | FBgn0087012 | Serotonin receptor 2A |
| 5-HT2B | FBgn0261929 | Serotonin receptor 2B |
| 5-HT7 | FBgn0004573 | Serotonin receptor 7 |
| Oct-TyrR | FBgn0004514 | Octopamine/tyramine receptor |
| Oamb | FBgn0024944 | Octopamine receptor in mushroom bodies |
| TyrR | FBgn0038542 | Tyramine receptor |
| norpA | FBgn0262738 | Phospholipase C (neurotransmission) |
| nSyb | FBgn0013342 | Synaptic vesicle protein |
| CSP | FBgn0004179 | Synaptic chaperone (Cysteine String Protein) |
| Shaker | FBgn0003380 | Voltage-gated potassium channel |
| para | FBgn0285944 | Voltage-gated sodium channel |
| cac | FBgn0263111 | Voltage-gated calcium channel |
| ATPsyn-beta | FBgn0010217 | ATP synthase β subunit (mitochondrial) |
| ND-75 | FBgn0017566 | NADH dehydrogenase subunit 75 kDa |
| SdhA | FBgn0000181 | Succinate dehydrogenase subunit A |
| COX5A | FBgn0019624 | Cytochrome c oxidase subunit 5A |
| Mhc | FBgn0264695 | Myosin heavy chain (muscle function) |
| Act88F | FBgn0000047 | Actin 88F (muscle-specific actin) |
| TpnC41C | FBgn0013348 | Troponin C isoform (muscle) |
| Tm1 | FBgn0003721 | Tropomyosin 1 (muscle) |
| per | FBgn0003068 | Period (circadian rhythm) |
| tim | FBgn0014396 | Timeless (circadian rhythm) |
| clk | FBgn0023076 | Clock (circadian rhythm) |
| cyc | FBgn0023091 | Cycle (circadian rhythm) |
| Pdf | FBgn0015271 | Pigment dispersing factor (circadian output) |
| mt: ND3 | FBgn0013681 | mitochondrial NADH-ubiquinone oxidoreductase chain 3 |
